# Genome sequencing and physiological characterization of three *Neoarthrinium moseri* strains

**DOI:** 10.1101/2025.04.28.650913

**Authors:** Nadine J. Hochenegger, Gabriel A. Vignolle, Matthias Schmal, Robert L. Mach, Astrid R. Mach-Aigner, Mohammad Javad Rahimi, Chin Mei Chan, Feng M. Cai, Irina S. Druzhinina, Christian Zimmermann

## Abstract

Fungi play essential ecological roles and have been utilized by humans for diverse applications such as industrial enzyme production or as sources of bioactive compounds. Recent research has highlighted the *Amphisphaeriales* order (*Ascomycota*) as promising producers of secondary metabolites of pharmaceutical importance. Within this family, the recently established genus *Neoarthrinium* includes species such as *N. brasiliensis, N. lithocarpicola, N. moseri, N. trachycarpi,* and *N. urticae*. Existing literature has primarily focused on the taxonomy and phylogeny of *Neoarthrinium*, leaving its physiology, ecology, and metabolic potential unexplored. This study presents the first investigation of the metabolic and genomic potential of *N. moseri*. We describe the isolation of two South-Asian *N. moseri* strains and the genome sequencing of these strains alongside the Colombian ex-type strain for the species. Comparative genome analysis reveals an exceptionally high number of biosynthetic gene clusters (BGCs), surpassing those of many other fungi in the *Amphisphaeriales* order. Additionally, the genome of *N. moseri* contains a diverse repertoire of carbohydrate-active enzymes (CAZymes), supporting its hypothesized ecological role as a phyllosphere fungus (putatively an endophyte and/or saprotroph). Ecophysiological assays, including BIOLOG phenotyping, demonstrate its ability to utilize a broad range of carbon sources, emphasizing ecological versatility.

This study highlights *N. moseri* as a promising candidate for secondary metabolite discovery, providing foundational insights into the metabolic and genomic potential of the *Neoarthrinium* genus and related fungi.

## Introduction

Fungi are a diverse kingdom with a broad range of ecological roles, e.g. decomposers of organic matter and symbiotic partners of plants – may it be in mutualistic and parasitic relations. Humankind has been using fungi for different purposes, such as food and feed fermentation, agricultural applications, enzymes production, and as source of bioactive compounds for medicine and industry [1]. The fungal secondary metabolism is generally considered to be a large untapped reservoir for novel bioactive compounds and drug leads [2], [3]. The ongoing efforts to find new pharmaceuticals encompass the search for new fungi and mining their genomes [4], [5], [6].

In the recent years, the order *Amphisphaeriales* (*Ascomycota*) gained increasing attention as promising secondary metabolite producers [7], especially, fungi related to such genera as *Apiospora* and *Arthrinium* (in the family of *Apiosporaceae*) were [8], [9].

The genus *Neoarthrinium*, established in 2022 within the family *Apiosporaceae (vide infra)*, originally comprised four species: *N. lithocarpicola*, *N. moseri*, *N. trachycarpi*, and *N. urticae* [10]. These taxa were isolated from diverse terrestrial plant hosts across Asia and South America: *N. lithocarpicola* from diseased leaves of *Lithocarpus glaber* in China; *N. trachycarpi* from *Trachycarpus fortunei*, also in China; *N. moseri* from a dead plant in Colombia; and *N. urticae* from leaf litter in India. The reclassification of *N. urticae* (syn. *Arthrinium urticae*) was based on sequence data from a single isolate, and its taxonomic placement remains tentative due to uncertainties regarding its representativeness [10]. Since then, three additional species have been described. *N. brasiliensis*, added in 2024, further expanded the genus into South America [11]. In 2025, two ecologically and geographically distinct taxa were introduced: *N. lewisiae* was isolated from necrotic leaf spots on *Pandanus tectorius* (screwpine) in coastal Australia [12] and *N. aquaticum* was described from submerged plant tissue of the golden leather fern (*Acrostichum aureum*) in a freshwater habitat in Thailand [13], representing the first aquatic species in the genus. The later study also proposed the establishment of the new family *Neoarthriniaceae*, to accommodate the genus *Neoarthrinium* [13]. These findings highlight the ecological versatility of *Neoarthrinium*, encompassing both saprobic and potentially pathogenic lifestyles across tropical and subtropical regions, with substrates ranging from terrestrial leaf litter and petioles to aquatic ferns and coastal monocots.

However, W. Gams isolated the *N. moseri* type strain CBS 164.80 from the dead petiole of *Mauritia minor* in Colombia in 1995 [14]. It was originally described as an unusual species of the genus *Wardomyces* (Microascales), currently reclassified into four genera, but Jiang et al. realized that this isolate belongs to *Amphisphaeriales* and assigned it to *N. moseri* in 2022 [10]. Up till now, all *Neoarthrinium* strains have been isolated from the surface of plants [10], [11], [12], [13], [14], [15], [16]. Limited to mostly taxonomic and phylogenetic studies, existing literature provides little insight into the physiology, ecology, or metabolic potential of this genus.

This study provides the first investigation of the metabolic and genomic potential of a species within the genus *Neoarthrinium*, namely *N. moseri*. We describe the isolation of two new *N. moseri* strains from Borneo and genome sequencing of these strains and the Colombian ex-type strain CBS 164.80. Our results reveal that *N. moseri* possesses an outstandingly large array of biosynthetic gene clusters (BGCs), exceeding those of many other *Amphisphaeriales* fungi. Furthermore, genome analyses highlighted not only an abundance of BGCs but also a complement of carbohydrate-active enzymes (CAZymes), matching its supposed role as epiphyte or saprobe. We explored basic growth characteristics and substrate utilization of *N. moseri* to gain physiological insights. A BIOLOG Phenotype microarray revealed its ability to utilize a wide variety of carbon sources, underscoring its ecological versatility. Additionally, our genomic data together with already existing phylogenetics and a reassessment of the spore sizes strongly suggest that *N. trachycarpi* should not be considered a separate species but strains of *N. moseri*.

## Results

### Isolation and DNA Barcoding of two epiphytic N. moseri strains

In 2008, we isolated the two epiphytic fungi TUCIM 5799 and TUCIM 5827 from the adaxial surface of the healthy high canopy leaf of *Shorea johorensis* (Dipterocarpaceae, Malvales; DNA BarCode maturase K (*matK*) deposited in NCBI GenBank MF993320.1 [17]) on Borneo.

The two new isolates as well as the reference strain CBS 164.80 form a light-colored mycelium (beige on malt extract and Czapek yeast autolysate plates, white on oatmeal after 14 days, Fig. 1A-I). The texture of the mycelia and the size of the colonies differ amongst the strains and depend on the culture media. Further, we observed the formation of a large quantity of conidia on oatmeal plates. The conidia were present in a slimy layer on the surface of the colonies. The strain TUCIM 5799 also produced smaller amounts of conidia on malt extract and Czapek yeast autolysate plates. The conidia of the two new isolates and the *N. moseri* reference strain look alike: They are melanized, dark colored, and are most frequently pear-shaped, with a length of 4.2 – 6.7 µm long and a width of 3 – 3.9 µm, in contrast to essentially larger dimensions previously reported for this species [14] (Fig. 1J-O, Table 1) and more consistent with the values obtained for *N. trachycarpi* (6.1–8.5 × 4.2–5.3 μm) [15]. Thus, we re-assessed Gams’ SEM images [14], and measured actually similar sizes we obtained from our own picture (Table 1).

**Figure 1.**
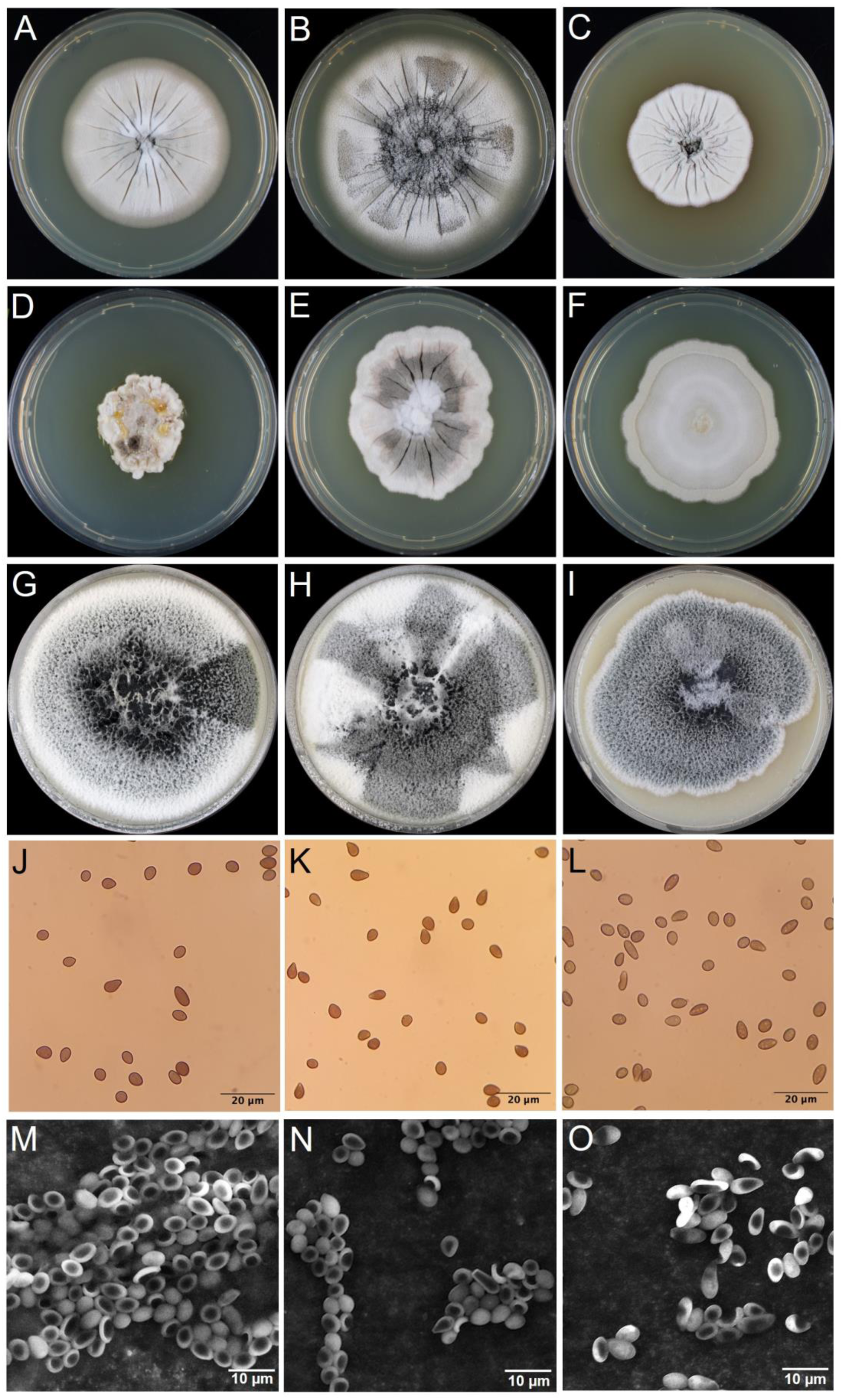
Morphology of *N. moseri* CBS 164.80 (A, D, G), TUCIM 5799 (B, E, H), and TUCIM 5827 (C, F, I) on malt extract (A-C), Czapek yeast autolysate (D-F), and oatmeal (G-I) plates after incubation at 28 °C for 14 days. Brightfield and scanning electron microscopy of spores of *N. moseri* CBS. 164.80 (J, M), TUCIM 5799 (K, N), and TUCIM 5827 (L, O).

**Table 1.** Average conidia dimensions.

| Strain | Length [µm] | Width [µm] | Reference |
| --- | --- | --- | --- |
| *N. moseri* CBS 164.80 | 10 - 14 | 3 – 4.5 | [14] (original values) |
| *N. moseri* CBS 164.80 | 4.5 – 7.1 | 3.1 – 4.3 | Re-assessment of pictures from [14] |
| *N. moseri* CBS 164.80 | 4.4 - 6 | 3.1 – 4.1 | This publication |
| *N. moseri* TUCIM 5799 | 4.2 – 5.7 | 3.2 – 3.8 | This publication |
| *N. moseri* TUCIM 5827 | 4.3 – 6.7 | 3 – 3.9 | This publication |
| *N. trachycarpi* | 6.1 - 8.5 | 4.2 - 5.8 | [15] |
| *N. lithocarpicola* | 5 – 8.5 | 4.5 - 6 | [10] |
| *N. brasiliensis* | 4 - 5 | 3 - 4 | [11] |
| *N. aquaticum* | 8 – 13 | 6 - 10 | [13] |
| *N. urticae* | n/a | n/a |  |
| *N. lewisiae* | n/a | n/a |  |

To confirm the classification of the new isolates as *N. moseri* strains, we performed a multiple sequence alignment of the available ITS, LSU and *tub2* sequences of the *Neoarthrinium* strains and constructed a phylogenetic tree (Fig. 2). *N. moseri* CBS 164.80 clusters together with two *N. trachycarpi* strains, *N. urticae* and the two new isolates.

**Figure 2.**
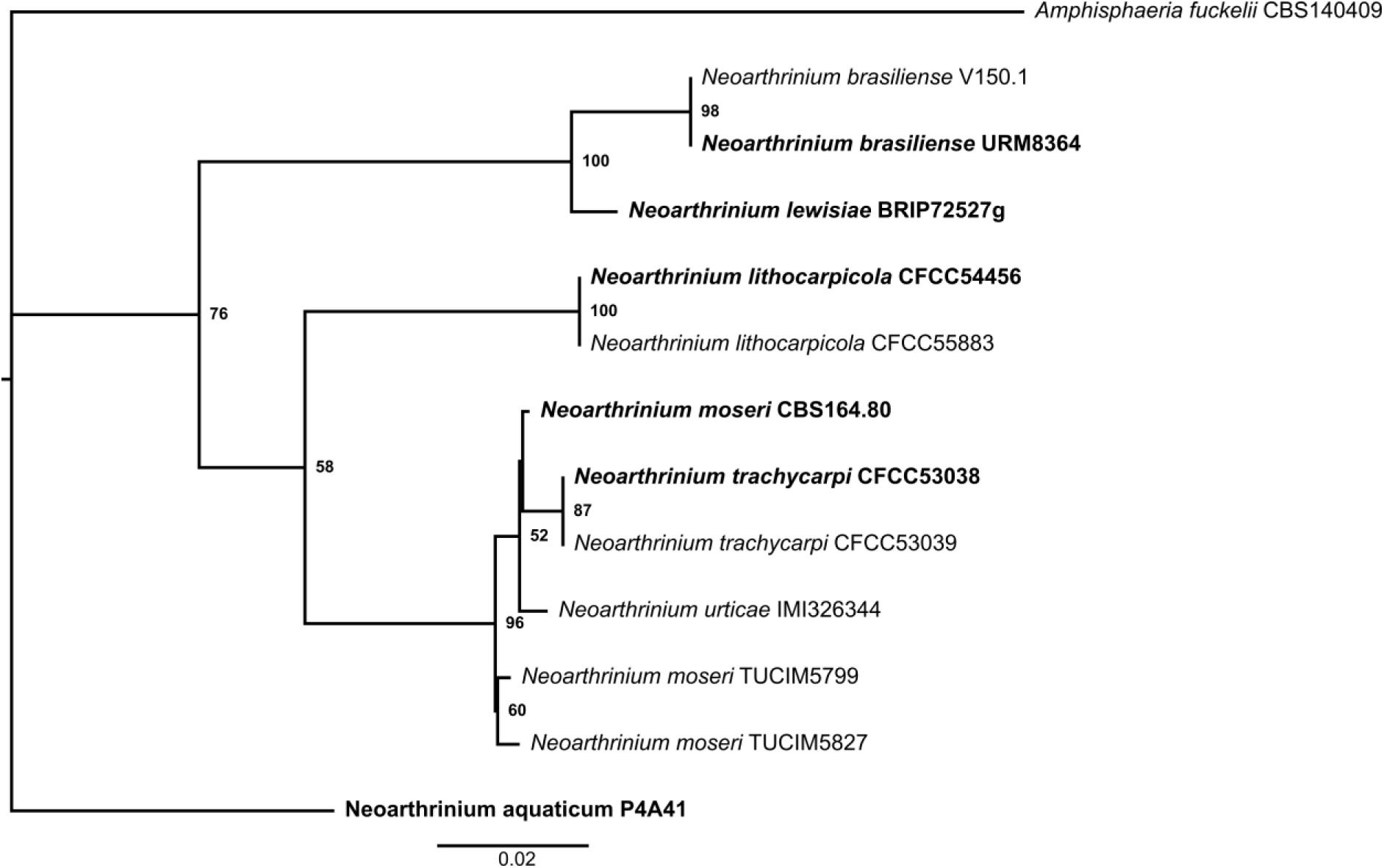
Phylogenetic tree based on the concatenated multiple sequence alignment of ITS, LSU and *tub2* of the indicated fungal isolates (type strains in bold). The rooted phylogenetic tree is the consensus of 10 individual runs applying 1000 bootstraps utilizing the maximum-likelihood approach. Values at nodes indicate bootstrap support values in %.

### Mitochondrial and nuclear genome of N. moseri

The extracted circularized mitochondrial genomes have a length of 42,769 bp, 43,978 bp, and 42,769 bp and with GC contents of 27.52%, 27.52%, and 27.53% for the strain CBS 164.80, TUCIM 5799, and TUCIM 5827, respectively (Fig. S1). The respective average sequencing coverages were at 364x, 8,939x, and 464x.

The size of *N. moseri* nuclear genomes is between 43.7 Mbp and 46.1 Mbp with average sequencing coverages between 32x and 141x. The detailed results of the genomes and assembly characteristics (size, GC content, characteristics for scaffold number and size, N_50_ and L_50_) are summarized in Table S1. To evaluate the completeness of the genome assembly, we performed a Benchmarking Universal Single-Copy Orthologues (BUSCO) analysis with the eukaryote dataset [18]. 100% complete BUSCOs without duplicates were found in all three assemblies (Table S1). Further, we calculated the average nucleotide identity (ANI) and found the three strains to be highly similar (Table S2). Additionally, the genomes of the three strains exhibited a similar GC content of around 52.7%, and masked element analysis indicated a low level of repetitive sequences, with simple repeats and low complexity regions occupying less than 1% of the genomes.

### Gene prediction and annotation

First, we identified and masked the repetitive elements in the nuclear genomes of the three *N. moseri* strains (Table 2). Additionally, we performed an tRNA prediction and found a total of 196, 190 and 189 tRNA genes, respectively (Table 2, Additional files 1-3).

**Table 2.** Masked repetitive elements and tRNA genes found in the genomes of the *N. moseri* strains CBS 164.80 (CBS), TUCIM 5799 (5799) and TUCIM 5827 (5827).

| Masked element | Number of elements* |  |  | Length occupied in bp |  |  | Percentage of sequence |  |  |
| --- | --- | --- | --- | --- | --- | --- | --- | --- | --- |
| Strain | CBS | 5799 | 5827 | CBS | 5799 | 5827 | CBS | 5799 | 5827 |
| SINEs | 35 | 35 | 33 | 2,289 | 2,404 | 2,231 | 0.01% | 0.01% | - |
| LINEs | 223 | 222 | 220 | 16,838 | 17,399 | 16,757 | 0.04% | 0.04% | 0.04% |
| LTR elements | 4 | 3 | 3 | 300 | 204 | 200 | - | - | - |
| DNA elements | 50 | 49 | 55 | 3,751 | 3,598 | 4,260 | 0.01% | 0.01% | 0.01% |
| Unclassified | 1 | 1 | 1 | 142 | 142 | 72 | - | - | - |
| Small RNA | 86 | 78 | 74 | 12,181 | 12,157 | 11,815 | 0.05% | 0.03% | 0.05% |
| Simple repeats | 7,572 | 7,432 | 7,532 | 306,538 | 294,925 | 297,695 | 0.70% | 0.66% | 0.64% |
| Low complexity | 652 | 600 | 652 | 30,991 | 27,670 | 30,995 | 0.06% | 0.06% | 0.07% |
| tRNA | 196 | 189 | 190 | 17,154 | 16,788 | 16,884 | 0.04% | 0.04% | 0.04% |

As no transcriptome data were available, the gene prediction was performed on the masked genome using a model trained with the genome of *Pestalotiopsis fici*. We obtained approx. 14,000 genes for *N. moseri* (Table 3). A significant portion of the predicted genes (34.4% to 36.7%) did not match any sequences in public databases below the E^-5^ threshold (Table 3), suggesting the presence of potentially novel genes unique to *N. moseri*.

**Table 3.** Gene predictions.

| Strain | Predicted putative genes | genes without BLAST hits below E<sup>-5</sup> |
| --- | --- | --- |
| CBS 164.80 | 13,929 | 4,797 (34.4%) |
| TUCIM 5799 | 14,160 | 4,964 (35.0%) |
| TUCIM 5827 | 14,595 | 5,352 (36.7%) |

The predicted gene sets were annotated by blasting them against the UniProt database and via the PANNZER2 web interface. The combined functional annotations are given in Additional Files 4-6.

### Genome mining for CAZymes

The ability to decompose organic matter and the saprotrophic lifestyle are hallmarks of fungal biology. Fungi thrive on plant biomass and other natural materials by degrading complex and simple carbohydrates using so-called carbohydrate active enzymes (CAZymes) [19]. We used dbCAN2 (a meta-server for CAZyme annotation) and a HMMer (Hidden Marcov model) search [20], a DIAMOND search [21], and a Hotpep search [22] to predict the CAZymes in the three *N. moseri* genomes (Fig. 3, Table 4). In total, 1,005, 1,011, and 1,018 CAZymes were predicted by all three methods in CBS 164.80, TUCIM 5799, and TUCIM 5827, respectively (Fig. 3, Additional files 7-9, including 455, 455, and 460 genes predicted by all three methods (Fig. 3).

**Figure 3.**
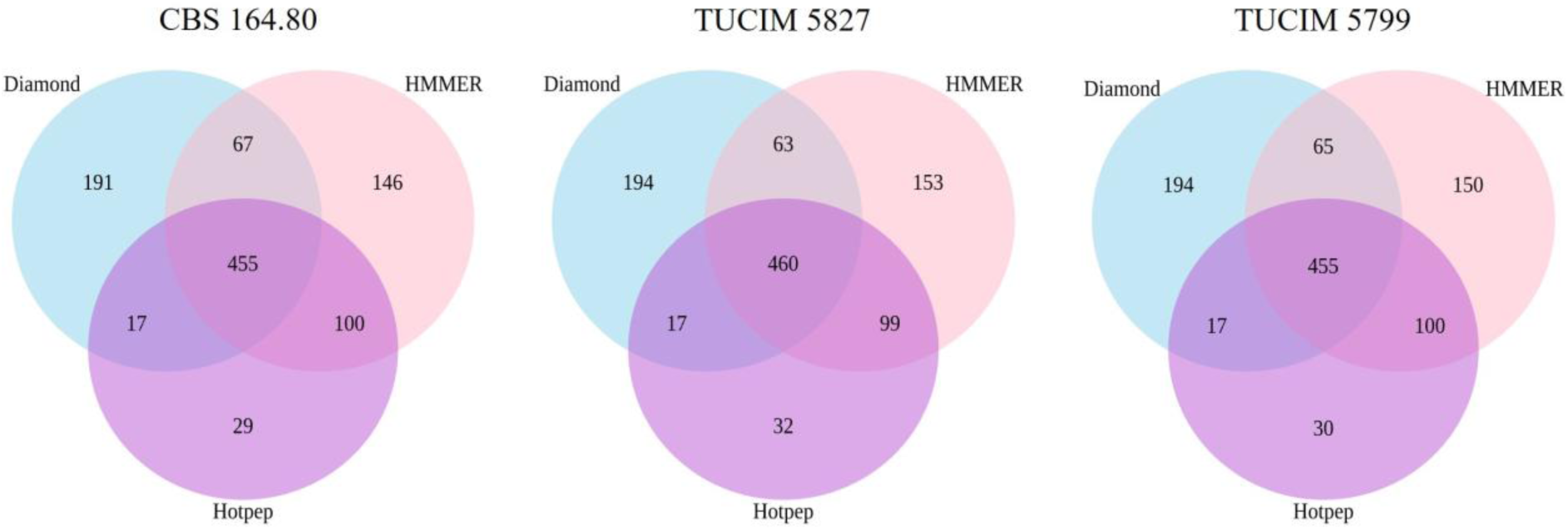
The genomes of the three sequences N. moseri strains were mined for putative CaZymes using Diamond, HMMER, and Hotpep.

**Table 4.** The carbohydrate active enzymes (CAZymes) found with dbCAN2 a meta-server for CAZyme annotation. Glycosyltransferases (GT); Glycoside Hydrolases (GH); carbohydrate esterases (CE); polysaccharide lyases (PL); Redox enzymes with auxiliary activities (AA).

| Strain | Total | GT | GH | CE | PL | AA |
| --- | --- | --- | --- | --- | --- | --- |
| CBS 164.80 | 1005 | 148 | 476 | 93 | 27 | 222 |
| TUCIM 5799 | 1011 | 152 | 479 | 94 | 27 | 222 |
| TUCIM 5827 | 1018 | 151 | 476 | 95 | 27 | 231 |
| *Pestalotiopsis fici* | - | 121 | 460 | 138 | 39 | - |

The dbCAN2 server also predicts certain subclasses of CAZyme. Glycosyltransferases (GT families) catalyze glycosidic bond formation and inversion and are part of the posttranslational modification steps in different compound formation processes. Glycoside hydrolases (GH families) is a large group of enzyme families which hydrolyse glycosidic bonds. Carbohydrate esterases (e.g., CE1, CE10 families) catalyze de-N or de-O-acylation of ester bonds in saccharides like in pectin. Polysaccharide lyases (e.g., PL1, PL7 families) cleave polysaccharide chains via β-elimination. Redox enzymes with auxiliary activities are involved in the breakdown processes of polysaccharides and lignin. The respective numbers of the predicted CAZymes subclasses (*sensu* dbCAN2) are also listed in Table 4. We used *Pestalotiopsis fici* as reference as it was the closest-related fungus with a complete genome sequenced.

### Genome mining for secondary metabolites

We used antiSMASH [23] to mine the genomes of the three *N. moseri* strains for genes that might be involved in the production of secondary metabolites and compared them to a few fungi of the same order (*Amphisphaeriales*) and the sister-taxon, the *Xylariales*, as well as the proficient secondary metabolite-producers *Aspergillus flavus*, *A. fumigatus* (*Eurotiales*) [24] and *Fusarium oxysporum* species complex (*Hypocreales*) [25]. Notably, the three *N. moseri* strains exhibited the highest BGC-count among the compared strains (85 in CBS 164.80, 88 in TUCIM 5799, 90 in TUCIM 5827) (Figure 4, Additional File 10).

**Figure 4.**
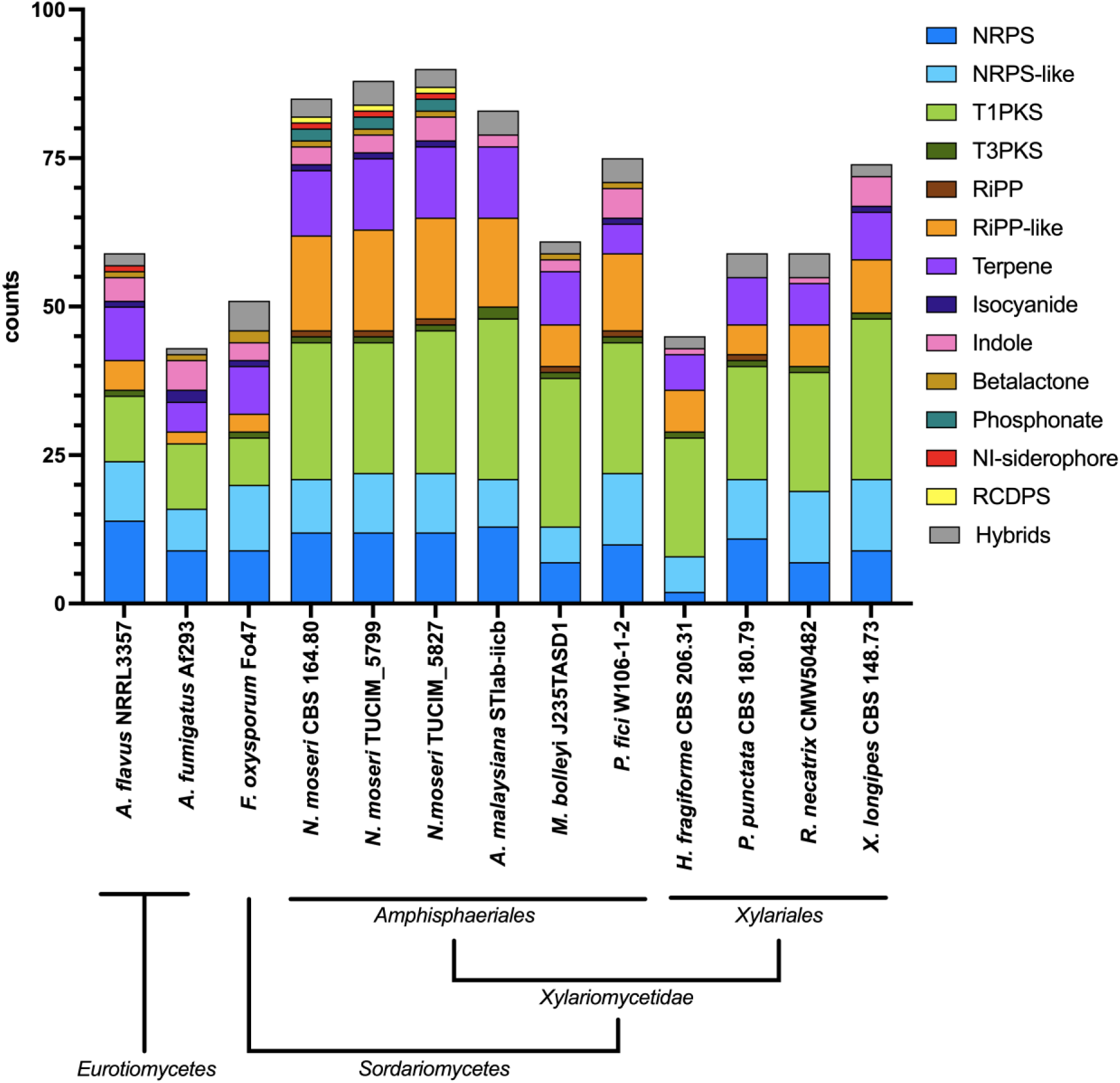
Overview of the predicted BGCs (antiSMASH 7.0) in the genomes of the indicated fungal species, including the three sequenced *N. moseri* strains (*Amphisphaeriales* genus *incertae sedis*). NRPS, non-ribosomal peptide synthetase; PKS, polyketide synthase; RiPP, ribosomally synthesized and post-translationally modified peptides; RCDP, arginine-containing cyclodipeptide synthase; hybrids, BGCs that contain core enzymes with characteristics for different classes, e.g. PKS-NRPS fusion enyzmes.

To get a better understanding of the secondary metabolite potential, we compared the predicted BGCs of the *N. moseri* strains to the MIBIG 4.0 database [26]. We assessed the predictions by a manual BGC comparison using the cblaster tool [27] if the BGC contained more than one gene (Additional File 11). We found 14 BGC similar to previously characterized BGCs (Additional File 12). Additional to BGCs for common compounds, such as siderophores or choline, we also found BGCs for antibacterial and antifungal substances, such as citridone A and related compounds, fusaric acid, and (-)-mellein. We also found BGCs highly similar to BGCs reported from plant pathogenic fungi, e.g. brassicicene C and koraiol, and the plant growth hormone gibberellin. Further, we detected BGCs for pharmaceutically interesting compounds such as the histone deacetylase inhibitor depudecin, the immunomodulator swainsonine, and the cytotoxin eupenifeldin.

### Growth optima and stress tolerance

To gain some insights into the ecophysiology of *N. moseri,* we cultivated the three strains on malt extract plates (MEX) with varying NaCl concentrations, pH values, or at different temperatures (Fig. 5A-C, individual growth curves are depicted in Fig. S2). All three *N. moseri* strains grew on MEX with NaCl concentrations ranging from 0 to 2.5 M (Fig. 5A), with optimal growth at 0 M and the least at 2.5 M. Mycelial growth progressively decreased as NaCl concentration increased.

**Figure 5.**
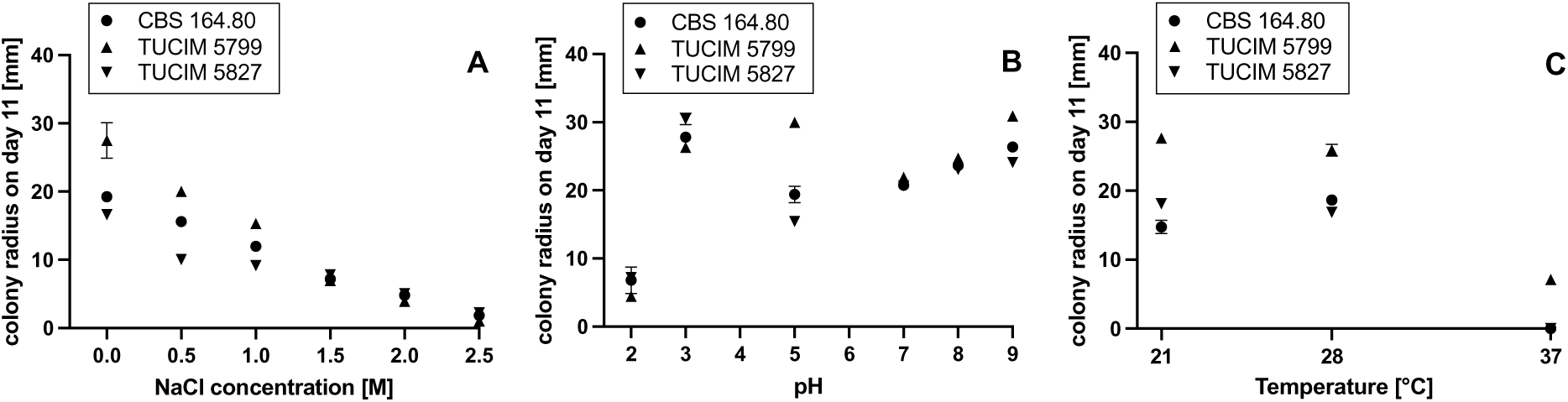
Growth response of the *N. moseri* strains NaCl-concentrations (A), pH (B), and temperatures (C), respectively, after 11 days of incubation on MEX. Data shows the mean of three experiments ± SD.

All three *N. moseri* strains grew on the media adjusted to any of the pH values between 2 and 9 (Fig. 5B), with pH 2 as the least tolerated condition. All three strains grew comparatively well at pH 3 with CBS 164.80 and TUCIM 5827 having their optima at this condition. In contrast, TUCIM 5977 has its growth optimum at pH 5, which was the second-least favorite condition for the other two strains. Regarding pH 7, 8, and 9, the three strains behaved similar to each other and showed modest growth. The optimum in the alkaline pH range was surprisingly at pH 9.

All three *N. moseri* strains grew at 21 °C and 28 °C (Fig. 5C). There was no difference in growth of TUCIM 5799 and TUCIM 5827 between these incubation temperatures. CBS 164.80 grew slightly better at 28 °C than at 21 °C. Only TUCIM 5799 was able to tolerate 37 °C but showed poor growth.

### Utilization of different carbon sources

We performed a BIOLOG Phenotype microarray to assess the carbon source utilization profiles of the three *N. moseri* strains. The growth was monitored over 16 days, and we used the maximal produced biomass throughout the growth period (OD_750_max) for comparison (Fig. 6, Table 5). The individual growth curves on each carbon source are shown in the supplements (Fig. S3-S10). The carbon sources were grouped as described previously [28]. “No growth” indicates OD_750_max values less than or equal to the respective OD_750_max on water. Neither of the three *N. moseri* strains grew on sedoheptulosan, L-sorbose, glucuronamide, N-acetyl-D-mannosamine, and 2-amino ethanol. TUCIM 5827 didn’t grow on N-acetyl-galactosamine and α-methyl-D-glucoside, either. On all other tested carbon sources we observed differently strong growth, suggesting that *N. moseri* has an adaptive and diverse primary metabolism.

**Figure 6.**
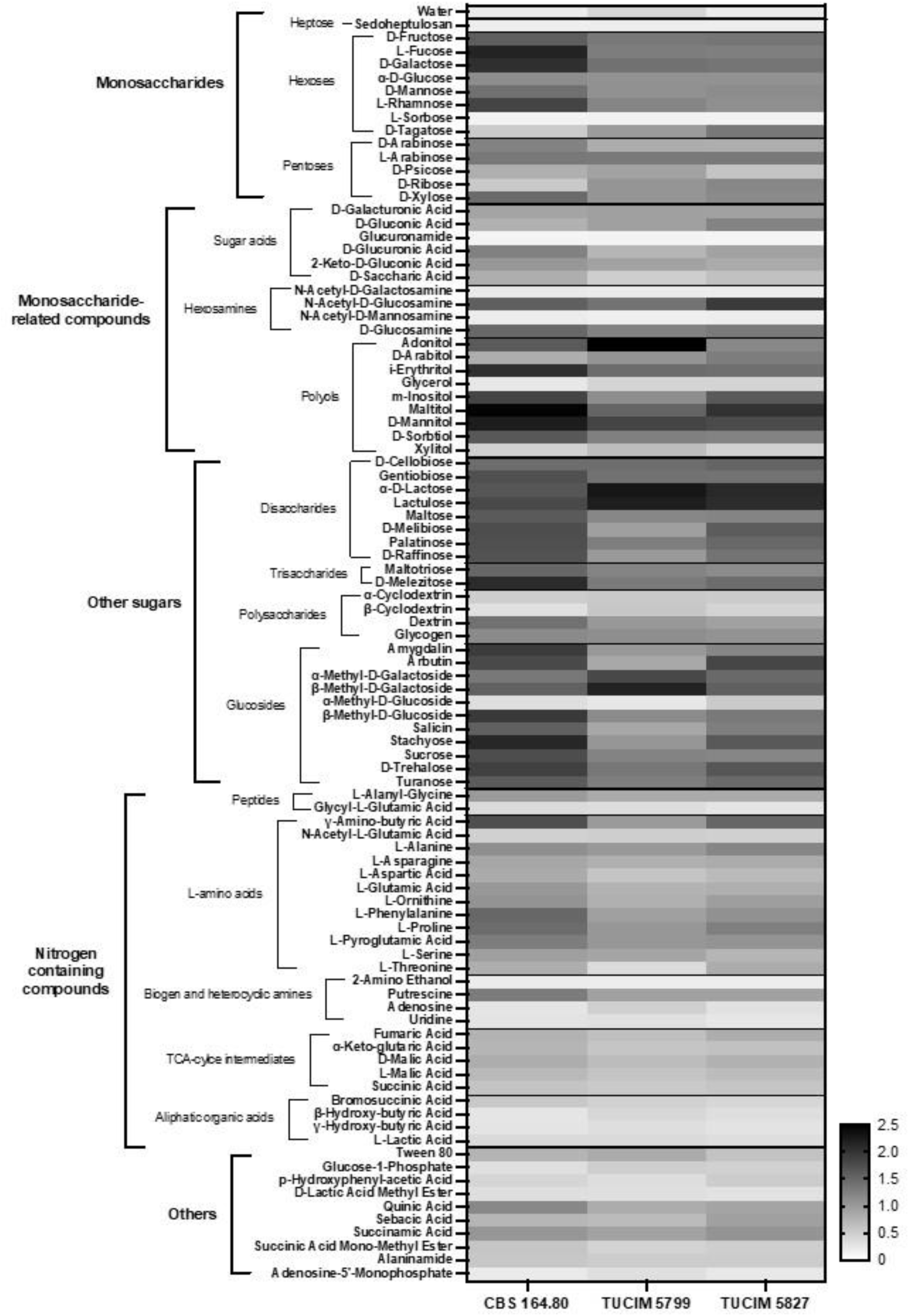
Comparative heatmap of the OD_750_max of the three tested *N. moseri* strains on the indicated carbon sources in BIOLOG FF microplates. Grayscale indicates respective OD_750_max.

**Table 5.** Growth of the *N. moseri* strains on different carbon sources. Strong growth was defined as an OD_750_max of 15% above the median OD_750_max for each strain, weak growth as 15% below the median OD_750_max, medium in between these two thresholds.

| Carbon-Source | CBS 164.80 | TUCIM_5799 | TUCIM_5827 | Carbon-Source | CBS 164.80 | TUCIM_5799 | TUCIM_5827 |
| --- | --- | --- | --- | --- | --- | --- | --- |
| Water | no |  |  | Monosaccharide-related compounds: Sugar acids |  |  |  |
| Monosaccharides: Heptose |  |  |  | D-Galacturonic Acid | medium |  |  |
| Sedoheptulosan | no |  |  | D-Gluconic Acid | medium |  |  |
| Monosaccharides: Hexoses |  |  |  | Glucuronamide | no |  |  |
| D-Fructose | strong | medium | strong | D-Glucuronic Acid | medium |  |  |
| L-Fucose | strong |  | medium | 2-Keto-D-Gluconic Acid | medium |  |  |
| D-Galactose | strong |  |  | D-Saccharic Acid | medium | slight |  |
| α-D-Glucose | medium |  |  | Monosaccharide-related compounds: Hexosamines |  |  |  |
| D-Mannose | medium |  |  | N-Acetyl-D-Galactosamine | slight |  | no |
| L-Rhamnose | strong |  | medium | N-Acetyl-D-Glucosamine | strong |  |  |
| L-Sorbose | no |  |  | N-Acetyl-D-Mannosamine | no |  |  |
| D-Tagatose | slight | medium | strong | D-Glucosamine | strong |  |  |
| Monosaccharides: Pentoses |  |  |  | Monosaccharide-related compounds: Polyols |  |  |  |
| D-Arabinose | medium |  |  | Adonitol | strong |  | medium |
| L-Arabinose | medium | strong |  | D-Arabitol | medium |  | strong |
| D-Psicose | medium |  | slight | i-Erythritol | strong |  |  |
| D-Ribose | slight | medium |  | Glycerol | slight |  |  |
| D-Xylose | strong | medium |  | m-Inositol | strong | medium | strong |
|  |  |  |  | Maltitol | strong |  |  |
|  |  |  |  | D-Mannitol | strong |  |  |
|  |  |  |  | D-Sorbitol | strong |  | medium |
|  |  |  |  | Xylitol | slight | medium | slight |

**Table 5** Growth of the *N. moseri* strains on different carbon sources (continued)
| Carbon-Source | CBS 164.80 | TUCIM_5799 | TUCIM_5827 | Carbon-Source | CBS 164.80 | TUCIM_5799 | TUCIM_5827 |
| --- | --- | --- | --- | --- | --- | --- | --- |
| Other sugars: Disaccharides |  |  |  | Other sugars: Glucosides |  |  |  |
| D-Cellobiose | strong |  |  | Amygdalin | strong | medium |  |
| Gentiobiose | strong |  |  | Arbutin | strong | medium | strong |
| α-D-Lactose | strong |  |  | α-Methyl-D-Galactoside | medium | strong |  |
| Lactulose | strong |  |  | β-Methyl-D-Galactoside | strong |  |  |
| Maltose | strong |  | medium | α-Methyl-D-Glucoside | slight |  | no |
| D-Melibiose | strong | medium | strong | β-Methyl-D-Glucoside | strong |  |  |
| Palatinose | strong |  |  | Salicin | strong | medium | strong |
| D-Raffinose | strong | medium | strong | Stachyose | strong | medium | strong |
| Other sugars: Trisaccharides |  |  |  | Sucrose | strong |  | medium |
| Maltotriose | strong |  | medium | D-Trehalose | strong |  |  |
| D-Melezitose | strong |  |  | Turanose | strong |  |  |
| Other Sugars: Polysaccharides |  |  |  | Nitrogen containing compounds: Peptides |  |  |  |
| α-Cyclodextrin | slight |  |  | L-Alanyl-Glycine | medium |  |  |
| β-Cyclodextrin | slight | medium | slight | Glycyl-L-Glutamic Acid | slight |  |  |
| Dextrin | medium |  |  |  |  |  |  |
| Glycogen | medium |  |  |  |  |  |  |

**Table 5** Growth of the *N. moseri* strains on different carbon sources (continued)
| Carbon-Source | CBS 164.80 | TUCIM_5799 | TUCIM_5827 | Carbon-Source | CBS 164.80 | TUCIM_5799 | TUCIM_5827 |
| --- | --- | --- | --- | --- | --- | --- | --- |
| Nitrogen containing compounds: L-amino acids |  |  |  | Nitrogen containing compounds: TCA-Cylce Intermediates |  |  |  |
| γ-Amino-butyric Acid | strong | medium | strong | Fumaric Acid | medium |  |  |
| N-Acetyl-L-Glutamic Acid | slight |  |  | α-Keto-Glutaric Acid | medium |  | slight |
| L-Alanine | medium |  |  | D-Malic Acid | medium |  |  |
| L-Asparagine | medium |  |  | L-Malic Acid | medium |  |  |
| L-Aspartic Acid | medium |  |  | Succinic Acid | slight |  |  |
| L-Glutamic Acid | medium |  |  | Nitrogen containing compounds: Aliphatic Organic Acids |  |  |  |
| L-Ornithine | medium |  |  | Bromosuccinic Acid | slight |  |  |
| L-Phenylalanine | strong | medium |  | β-Hydroxy-butyric Acid | slight |  |  |
| L-Proline | strong | medium |  | γ-Hydroxy-butyric Acid | slight |  |  |
| L-Pyroglutamic Acid | medium |  |  | L-Lactic Acid | slight |  |  |
| L-Serine | medium |  |  | Others |  |  |  |
| L-Threonine | medium | slight | medium | Tween 80 | medium |  | slight |
| Nitrogen containing compounds: Biogene and Heterocyclic Amines |  |  |  | Glucose-1-Phosphate | slight |  |  |
| 2-Amino Ethanol | no |  |  | p-Hydroxyphenyl-acetic Acid | slight |  |  |
| Putrescine | medium |  |  | D-Lactic Acid Methyl Ester | slight |  |  |
| Adenosine | slight |  |  | Quinic Acid | medium |  |  |
| Uridine | slight |  |  | Sebacic Acid | medium |  |  |
|  |  |  |  | Succinamic Acid | medium |  |  |
|  |  |  |  | Succinic Acid Mono-Methyl Ester | slight |  |  |
|  |  |  |  | Alaninamide | slight |  |  |
|  |  |  |  | Adenosine-5'-Monophosphate | slight |  |  |

All three *N. moseri* strains showed high growth on saccharides (Fig. 6). In contrast, nitrogen-containing compounds were poorly utilized. For CBS 164.80, we observed strong growth only on γ-amino-butyric acid, L-phenylalanine, and L-proline. TUCIM 5827 grew well only on γ-amino-butyric acid and TUCIM 5799 did not metabolize any of the tested nitrogen-containing compounds well (Table 5).

## Discussion

In this study, we isolated and characterized two new strains of *N. moseri* (TUCIM 5799 and TUCIM 5827). Morphological and genetic analyses confirmed the attribution of these strains to *N. moseri*, while also revealing strain-specific variations in growth and morphological characteristics. *N. moseri* exhibited high adaptability across a range of growth conditions, including varying pH levels and salt concentrations, and demonstrated a diverse and adaptable carbon utilization pattern. Genome mining further uncovered an exceptional number of BGCs, highlighting the species’ considerable secondary metabolism potential. Additionally, an in-depth analysis of CAZymes revealed a rich repertoire consistent with a saprotrophic lifestyle, suggesting *N. moseri* plays a significant role in decomposing plant materials in its natural environment.

### Phylogeny and Taxonomic Insights

The type strain of *N. moseri* (CBS 164.80) was isolated in Colombia in 1995 and originally described as an unusual *Wardomyces* [14]. In 2022, Jiang et al. created the genus *Neoarthrinium* upon isolation of several new fungi in China [10].

Importantly, *N. moseri*, *N. trachycarpi*, and *N. urticae* possess highly similar ITS, LSU, and *tub2* sequences (Fig. 2), as already discussed by Jiang et al. and Mukhopadhyay *et al*. [10], [13]. However, Jiang *et al*. proposed keeping *N. moseri* and *N. trachycarpi* as separate species based on conidial size differences. Importantly, they compared their own measurements of *N. trachycarpi* to the values of *N. moseri* as reported by W. Gams. Our reassessment of Gams’ SEM images [14], along with our own conidial size measurements of the CBS strain and our two new isolates, revealed inconsistencies with previously reported values. It appears the spores of *N. moseri* and *N. trachycarpi* are similar in size (Table 1), suggesting that *N. trachycarpi* is not a distinct species. Further, the high ANI values among the CBS strain and our isolates (Table S2) strongly suggest that they belong to the same species. This fact, taken together with the phylogenetic tree based on the ITS, LSU, and *tub2* sequences (Fig. 2) and the analysis by Mukhopadhyay *et al*. [13] suggest that *N. moseri*, *N. trachycarpi*, and *N. urticae* belong to the same species.

Notably, the high ANI values among the sequenced *N. moseri* strains also indicate a stable genomic architecture and strong relatedness across the strains, despite spatiotemporal differences in their isolation.

The available sequences for *N. urticae* are from a single isolate from leaf litter in India (IMI 326344) but not from isolates from the type host *Urtica dioica* L. (*Urticaceae*). Jiang et al. already stated that “Additional molecular studies on verified isolates from *Urtica* collected in Europe are necessary to reveal whether IMI 326344 represents true *N. urticae*. However, *N. urticae* appears to be very rare and we are unaware of any additional collections with the exception of the type.” [10]. We second this opinion. Resolving these taxonomic uncertainties will also require more comprehensive sampling, sequence datasets, including less-conserved regions or population genomic analyses.

### Adaptability and Physiological Insights

Although the three sequenced *N. moseri* strains are genetically similar, they displayed distinct morphological traits under varying culture conditions (Fig. 1) and slight differences regarding their growth condition tolerance (Fig. 5) and carbon source utilization (Fig. 6 and Table 5). However, all strains exhibited strong conidiation on oat medium, making it a reliable choice for conidia collection in future experiments.

The BIOLOG assays revealed that all strains preferentially metabolized sugars, though with noticeable differences in biomass production. TUCIM 5827 consistently showed lower biomass accumulation across all substrates (Fig. 6 and Table 5), correlating with its weaker growth on MEX and oat media (Figure 2). The wide range of metabolized carbon sources suggests a highly versatile and adaptive catabolic system, reinforcing the idea that *N. moseri* is a saprotroph with considerable ecological resilience.

We also observed that γ-aminobutyric acid (GABA) was efficiently metabolized by CBS 164.80 and TUCIM 5827, while TUCIM 5799 did so moderately. This is particularly intriguing as plants produce GABA in response to stress, including fungal infections [29], [30]. While GABA can inhibit the growth of certain plant pathogens [31], [32], the ability of *N. moseri* to utilize GABA may confer an ecological advantage and may be a potential adaptation to an endophytic or epiphytic lifestyle. Further studies are needed to determine whether *N. moseri* can withstand inhibitory GABA concentrations or if it exclusively utilizes it as a nutrient.

The strains also exhibited a certain degree of halotolerance, characterized by optimal growth at low NaCl concentrations and reduced growth at higher concentrations (Figure 3A). Acidotolerance was observed, with CBS 164.80 and TUCIM 5827 thriving at pH 3, while TUCIM 5799 preferred pH 5 (Figure 3B). Importantly, the strains demonstrated growth across a broad pH range including even very high pH values, indicating their ability to survive in diverse environmental conditions. Thermotolerance varied among the strains, with TUCIM 5799 being the only one able to grow at 37 °C (Figure 3C). This observation is significant given the shared habitat of TUCIM 5799 and TUCIM 5827, suggesting a localized adaptation or microevolutionary divergence. In general, *N. moseri* can be classified as a mesophilic species, with strain-specific physiological adaptations that are likely to contribute to its ecological success.

### Mitochondrial genome

The mitochondrial genome of *N. moseri* contains 14 conserved protein-coding genes (Fig. S1), as expected for fungi, with one exception: the *atp8* gene (encoding for the ATP synthase F0 subunit 8) is absent. This gene was presumably transferred to the nuclear genome, as a single gene encoding for a putative ATP synthase subunit can be found in each genome of the three strains (JN550g13373 in *N. moseri* CBS 164.80; JX265g13592 in *N. moseri* TUCIM 5799; JX266g13823 in *N. moseri* TUCIM 5827). Such gene transfers are well-documented in fungi and highlight evolutionary genomic plasticity. This phenomenon may influence mitochondrial function and warrant further investigation into its implications for energy metabolism and strain-specific adaptation.

### Genome mining and metabolic potential

The extensive CAZyme repertoire in *N. moseri* (Table 4) supports the notion of a saprotrophic lifestyle, and indicates metabolic flexibility, allowing *N. moseri* to utilize a variety of complex and simple carbon sources, a trait confirmed by BIOLOG analysis that suggests an oligotrophic lifestyle.

Genome mining for BGCs revealed a striking potential for secondary metabolite production, consistent with its classification in the *Amphisphaeriales* order [8], [9]. *N. moseri* demonstrates a biosynthetic potential comparable to or even exceeding that of fungi known for and studied partly due to their secondary metabolism, such as *Aspergillus* and *Fusarium*. In general, secondary metabolites can be useful under certain conditions and contribute to an organism’s fitness. We found BGCs for common metabolites such as choline, the siderophore dimethylcoprogen, and the protective DHN-melanin, which contribute to the basic fitness of *N. moseri,* Additionally, *N. moseri* also possess BGCs that are most likely responsible for the production of antimicrobial compounds, which can contribute to fitness in competitive situations. Interestingly, we also found BGCs likely to produce brassicicene C and koraiol, and the plant growth hormone gibberellin, suggesting that *N. moseri* might not only be an inert epiphyte, but might directly interact with its plant host. Importantly, for most of the predicted BGCs, we could not detect similar BGCs in the MIBIG 4.0 database, suggesting that further exploration of these BGCs could lead to the discovery of novel compounds with pharmaceutical or agricultural applications.

## Materials and Methods

### Sampling and strain purification

The epiphytic fungi TUCIM 5799 and TUCIM 5827 were isolated from the adaxial surface of the healthy leaf of *Rubroshorea johorensis* (*Dipterocarpaceae*, *Malvales*; DNA BarCode maturase K (*matK*) deposited in NCBI GenBank MF993320.1, [17]) sampled in the high canopy (40 – 60 m above ground) of the lowland tropical rain forest surrounding the Kuala Belalong Field Studies Center (KBFSC, 4°32’48.2”N 115°09’27.9”E) located in the Temburong District of Brunei Darussalam (Borneo). For this purpose, the adaxial surface of a freshly sampled leaf was scratched by the sterile electric toothbrush (2 min) in 25 ml of sterile water supplemented with Tween-20 (0.01%) in large sterile Petri plate (20 cm in diameter). The resulting suspension was collected in 50 ml falcons and centrifuged at 4⁰C for 15 min at 14 000 rpm. The resulting pellet was resuspended in 4 ml of sterile water and used for serial dilution and plating on potato dextrose agar (PDA, Carl Roth) supplemented with 200 mg/l of chloramphenicol. Young single spore fungal colonies were detected with the use of a stereo microscope and aseptically transferred to fresh PDA plates and cultivated at 28°C in darkness. Agar plugs with pure mature cultures were preserved in 40% glycerol and stored at −80°C in TU Wien Collection of Industrial Microorganisms (TUCIM).

### Maintenance and morphological characterization of strains

CBS 164.80, TUCIM 5827, and TUCIM 5799 were maintained on agar plates containing 30 g/l oatmeal (S-Budget, SPAR Österreichische Warendhandels-AG; shredded to Ø 0.25 mm) and 15 g/l agar.

Three different media were used to initially evaluate the morphology of the three strains: MEA (20 g/l malt extract, 15 g/l agar), CYAS (Czapek yeast autolysate agar with 50 g/l NaCl; 3 g/l NaNO_3_, 5 g/l yeast extract, 30 g/l sucrose, 1.3 g/l K_2_HPO_4_, 0.5 g/l KCl, 0.5 g MgSO_4_ • 7 H_2_O, 0.01 g FeSO_4_ • 5 H_2_O, 0.005 g CuSO_4_ • 5 H_2_O, 15 g/l agar), and Oat (as above). 5 µl of spore solution (8 g/l NaCl, 0.05%(v/v) Tween-80) with an OD_600_ of 3.0 were applied to the middle of the agar plates, which were subsequently incubated at 28 °C for 13-14 days.

### Microscopy

Brightfield microscopy (BF) was performed using VWR Microscope TR 500 (VWR International GmbH, Darmstadt, Germany).

Scanning electron microscopy (SEM) of conidia was performed using COXEM EM-30AX PLUS with a SPT-20 Sputter. For sample preparation, conidia of the respective strain were softly scratched off an overgrown oatmeal-plate with a cotton swab. Conidia were then carefully distributed over a silver stripe which was attached to the stage of the device. Further proceedings were done according to the manufacturer’s instructions. Pictures were processed using the device’s own software Nanostation 3.0.4. The sizes of the spores were measured using FIJI (ImageJ2, Version 2.9.0).

### DNA extraction and library preparation

The *N. moseri* strains were cultivated in malt extract medium at 28°C and 180 rpm for 10 days in an orbital shaking incubator. The biomass was filtered through miracloth (EMD Millipore Corp., Burlington, MA, USA), frozen in liquid nitrogen, lyophilized. For DNA extraction, the lyophilized biomass was disrupted using a Fast-Prep-24 (MP Biomedicals, Santa Ana/, CA, USA) with 0.37 g of glass beads Ø 0.1 mm, 0.25 g of glass beads Ø 1 mm, and a glass bead Ø 5 mm at 6 m/s for 30 sec. After the addition of 1 ml CTAB buffer (100 mM Tris.Cl, 20 mM EDTA, 1.4 M NaCl, 2 % (w/v) CTAB, pH = 8.0) and 4 µl β-mercaptoethanol, the samples were subjected to two further disruption treatments on the Fast-Prep-24 at 5 m/s for 30 sec and then incubated at 65°C for 20 min. The supernatant was extracted with phenol, chloroform, isoamylalcohol (25:24:1) followed by a chloroform extraction. The supernatant was treated with RNase A (Thermo Fisher Scientific, Inc., Waltham, MA, USA) according to the manufacturer’s instructions. Finally, the DNA was precipitated with ethanol and dissolved in 10 mM Tris.Cl (pH = 8.0)

The DNA was sheared in a Diagenode Bioruptor^®^ Pico (Diagenode s.a., Liège, Belgium) with the settings set to high and three cycles of 15 sec “on” and 60 sec “off”. The sheared DNA was purified using PCR purification kit (Thermo Fisher Scientific, Inc., Waltham, MA, USA) and then double side size selected with “NEBNext Ultra^TM^ sample purification beads” (New England Biolabs, Ipswich, MA, USA) for 800 bp fragments. The library preparation was performed following the protocol of “NEBNext^®^ Ultra^TM^ II DNA Library Kit with Purification Beads” and “NEBNext^®^ Multiplex Oligos for Illumina (Index Primer Set1 and Set2)” (New England Biolabs, Ipswich, MA, USA). The average size in bp of the library was measured with the fragment analyzer from Advanced Analytical Technologies using the Agilent dsDNA 915 Reagent Kit (35-5000bp) and analyzed with the PRO size software (Agilent Technologies, Santa Clara, California, USA). The exact DNA concentrations were measured with an “invitrogen^TM^ Qubit^TM^ fluorometer” in ng/µl (Thermo Fisher Scientific, Inc., Waltham, MA, USA) using a “Quant-iT^TM^ dsDNA BR Assay” kit (Thermo Fisher Scientific, Inc., Waltham, MA, USA). Specifically, two libraries were created with a DNA fragment length of 1293 ± 6 bp and 1136 ± 7 bp, the average

DNA concentrations were 34.93 ± 0.25 ng/μl and 10.63 ± 0.23 ng/μl, resulting in a 40.899 nM and a 14.178 nM library, respectively. The libraries were diluted to the appropriate 4 nM concentration for sequencing.

### Sequencing

The sequencing of the *N. moseri* library was performed on an Illumina MiSeq platform using two V3 Reagent Kit (600 cycles) and one V2 Nano Reagent Kit (500 cycles) following the standard protocol of Illumina sequencing protocol without adding PhiX control to the runs (Illumina, San Diego, California, USA), resulting in a total of 67,670,936 paired end-reads. The raw data were deposited at the Sequence Read Archive (SRA) under the accession SRR13570309, SRR13747339 and SRR13747338. The quality profiles and all further figures, if not specified otherwise, were visualized in R [33].

### Extracting the mitochondrial genome and cleaning the raw reads

First, a preliminary assembly was performed using SPAdes v3.13.1 [34] with default parameters for each strain separately. Mitochondrial sequences were identified in each strain by performing a sequence similarity analysis using BLAST [35] (non-redundant nucleotide database). Contigs ranging from 500 to 1000 bp were then used as seed input for NOVOplasty v3.7 [36] to extract the whole circularized mitochondrial genome of *N. moseri* CBS 164.80, TUCIM 5799 and TUCIM 5827. This was performed in an iterative manner. The mitochondrial genomes were visualized with CGViewer [37]. The mitochondrial genomes were annotated with the automated MITOS2 web pipeline. The mitochondrial genomes were deposited at GenBank with accession no. MW554918, MW660808, and MW660809.

Using the mitochondrial genomes of the strains as input an index was built with bowtie v1.2.2 [38], respectively, and the mitochondrial flagged reads were extracted using --un option from each raw reads file. The clean raw reads were then re-paired with Fastq-pair [39] to use paired end read assemblers.

### Whole - genome assembly

For each strain respectively, the raw cleaned paired end reads were quality trimmed using Trimmomatic [40] in the command line and specifying PE for paired end reads and ILLUMINACLIP:Adapter-PE.fa:2:30:10:2:keepBothReads LEADING:3 TRAILING:3 SLIDINGWINDOW:4:15 MINLEN:36 to ensure high quality adapter-free reads. Then the cleaned raw reads were assembled using SPAdes v3.13.1 [34], for each strain separately. Furthermore, the high quality trimmed cleaned paired end reads were used for scaffolding with SSPACE-Standard v3.0 in an iterative manner with following command line options -x 1 -m 50 -o 20 -k 8 -a 0.70 -n 30 -z 150 –b and –k 6. Ns introduced during the assemblies and the scaffolding, so called gaps, were closed with GapFiller v1-10 [41] using following commands -m 30 -o 6 -r 0.7 -n 10 -d 50 -t 10 -g 0 -i 5 -b.

The assemblies were further improved by using Pilon v1.21 [42] iteratively. We first indexed the assemblies with bwa [43], SAMtools v1.7 [44] and picard [45]. The high quality trimmed cleaned paired end reads were mapped to the matching indexed assemblies of the individual *N. moseri* strains with bwa. The reads were mapped and combined in one step. Next, we sorted and created bam files from the sam files using SAMtools. Together with the paired sequencing reads, these were used as input for Pilon to iteratively improve each genome.

The genome assemblies were deposited at GenBank the accession no. GCA_022829205.1, GCA_022829195.1, and GCA_022829225.1.

### Phylogenetic analysis

Sequences used for the multiple sequence alignment can be seen in Table 6. Multiple sequence alignment was performed by using MAFFT [46]. The alignments for each gene were manually curated and concatenated using MEGA [47]. Based on this MSA iqtree2 was used to generate phylogenetic trees using the “K2P+I” nt substitute model [48]. Trees were generated by applying 1000 bootstraps in 10 individual runs each. Numbers at nodes indicate bootstrap support values in %. The tree was visualized using figtree software [49].

**Table 6.**
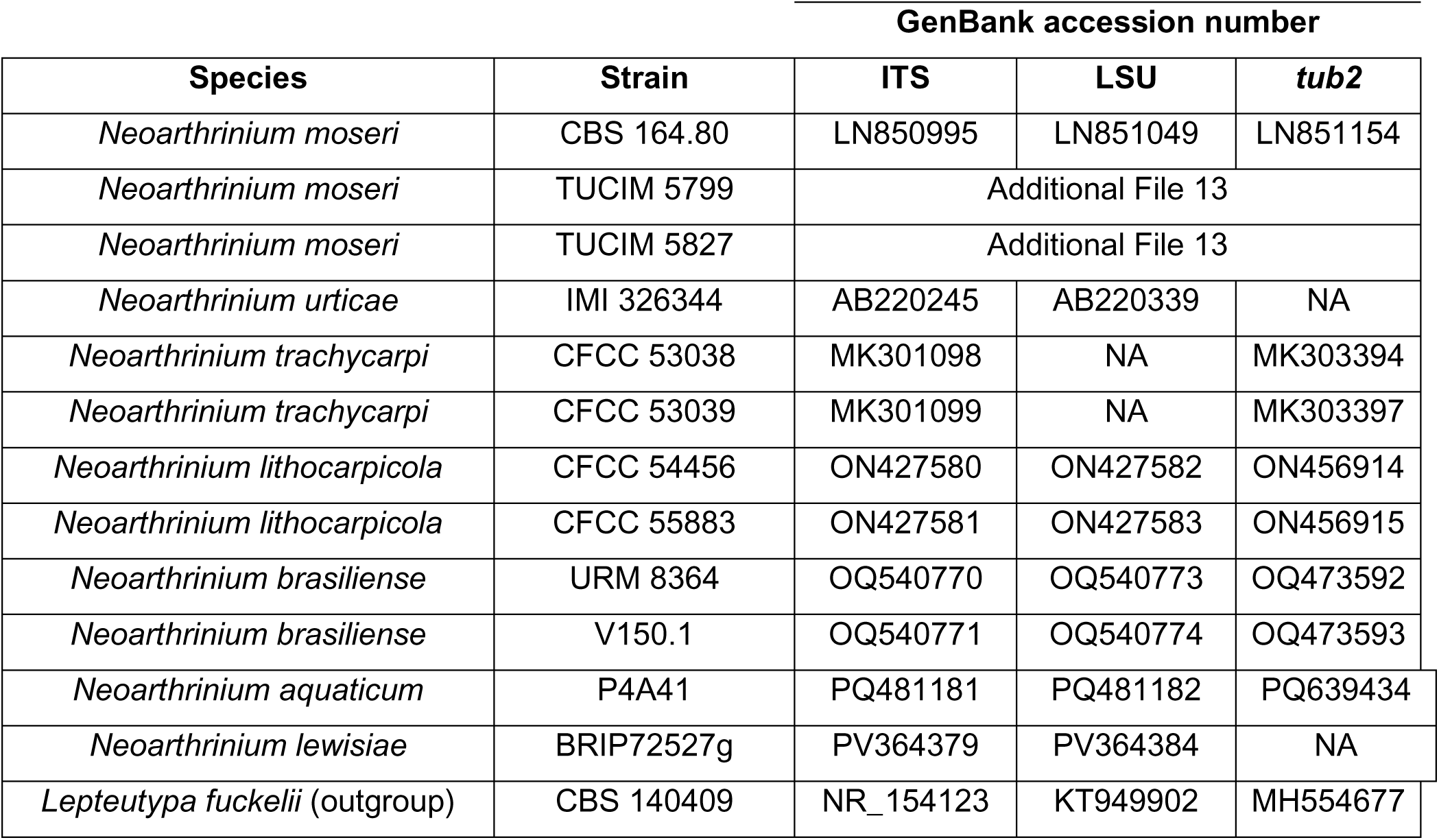
Isolates and GenBank accession numbers used in the phylogenetic analyses. NA, not available.

| Species | Strain | GenBank accession number |  |  |
| --- | --- | --- | --- | --- |
|  |  | ITS | LSU | *tub2* |
| *Neoarthrinium moseri* | CBS 164.80 | LN850995 | LN851049 | LN851154 |
| *Neoarthrinium moseri* | TUCIM 5799 | Additional File 13 |  |  |
| *Neoarthrinium moseri* | TUCIM 5827 | Additional File 13 |  |  |
| *Neoarthrinium urticae* | IMI 326344 | AB220245 | AB220339 | NA |
| *Neoarthrinium trachycarpi* | CFCC 53038 | MK301098 | NA | MK303394 |
| *Neoarthrinium trachycarpi* | CFCC 53039 | MK301099 | NA | MK303397 |
| *Neoarthrinium lithocarpicola* | CFCC 54456 | ON427580 | ON427582 | ON456914 |
| *Neoarthrinium lithocarpicola* | CFCC 55883 | ON427581 | ON427583 | ON456915 |
| *Neoarthrinium brasiliense* | URM 8364 | OQ540770 | OQ540773 | OQ473592 |
| *Neoarthrinium brasiliense* | V150.1 | OQ540771 | OQ540774 | OQ473593 |
| *Neoarthrinium aquaticum* | P4A41 | PQ481181 | PQ481182 | PQ639434 |
| *Neoarthrinium lewisiae* | BRIP72527g | PV364379 | PV364384 | NA |
| *Lepteutypa fuckelii* (outgroup) | CBS 140409 | NR_154123 | KT949902 | MH554677 |

**Table 7.**
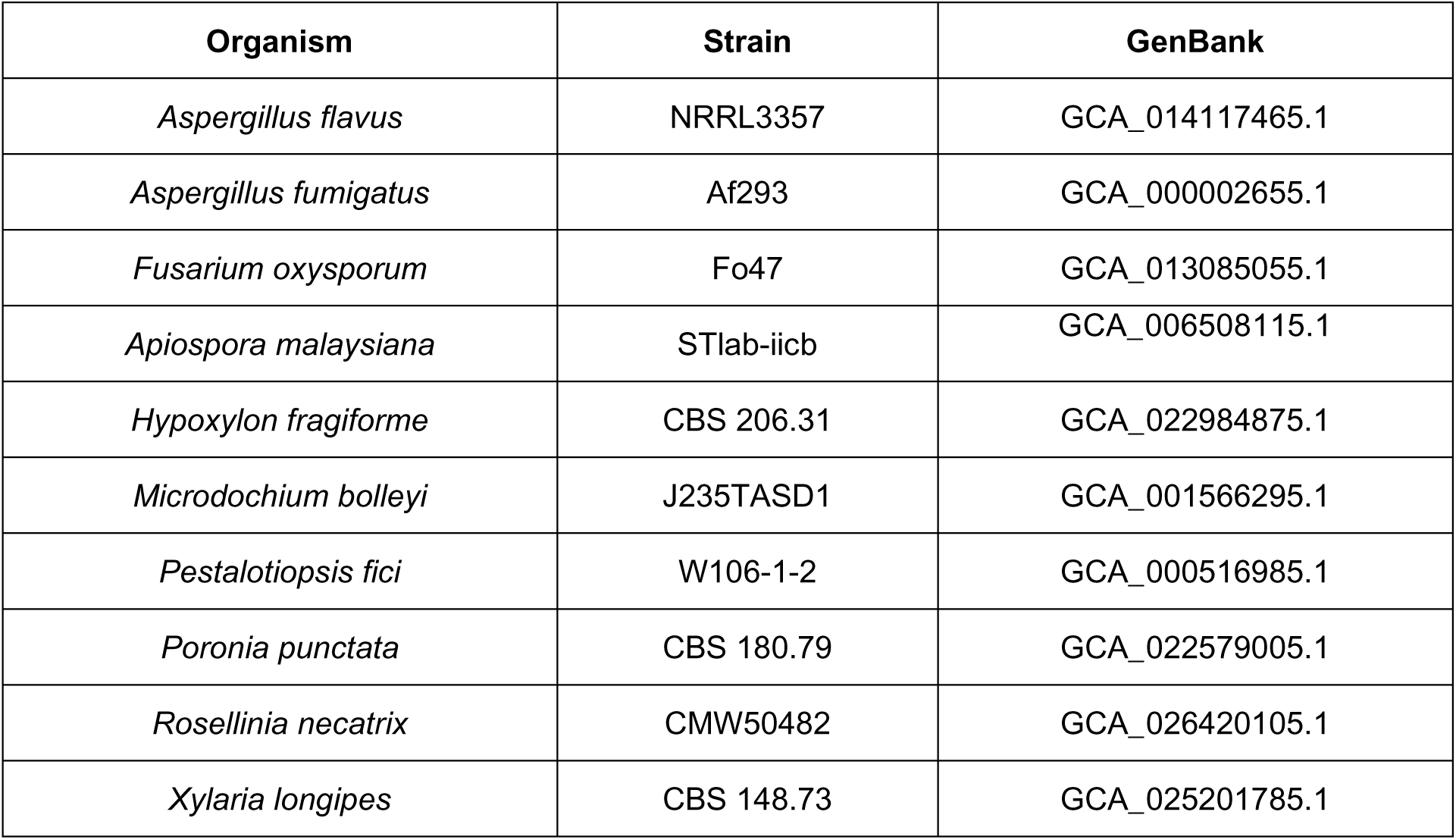
Genomes used for comparative antiSMASH genome mining.

### Gene prediction

To predict the genes, we first masked the repetitive elements in the nuclear genomes of *N. moseri* CBS 164.80 and our two new isolates to reduce the number of false positives during the subsequent gene prediction using RepeatMasker [50] Further, we performed an tRNA prediction with tRNAscan-SE v1.3.1 [51] using the unmasked genome. tRNAscan-SE.

For the gene prediction, we used Augustus v3.3.2 [44], because no transcriptome data was available. Augustus v3.3.2 [52] was trained with the genome of *Pestalotiopsis fici* (assembly PFICI; BioSample accession: SAMN02369365) following the protocol by Hoff & Stanke [53]. The genomes and the gene sets were evaluated using Quast v5.0.2 [54]. Quast v5.0.2 includes benchmarking with Benchmarking Universal Single-Copy Orthologs (BUSCO) v3.0.2, this was performed with the eukaryote dataset of 303 BUSCOs from 100 species. We further evaluated the gene predictions by aligning the amino acid sequences using Blastp v2.9.0+ [35] against the UniProt database [55].

### Annotation

The gene sets were first annotated using Blastp against the UniProt protein database. Protein ANNotation with Z-scoRE (PANNZER2) [56] was used to provide both GO and free text DE producing an accurate functional annotation.CAZymes were annotated using the dbCAN2 [57] meta server by applying a HMMer (Hidden Marcov model) search [20], a DIAMOND [21] search and a Hotpep [22] search and combining the three outputs.

### BGC Genome mining

The antiSMASH 7.1.0 fungal-version web version [23] was used for genome mining for secondary metabolite BGCs with following extra features applied: KnownClusterBlast, ClusterBlast, SubClusterBlast, MIBiG cluster comparison, ActiveSiteFinder, RREFinder, Cluster Pfam analysis, Pfam-based GO term annotation, TIGRFam analysis.

### Ecophysiological profiling

MEX-medium (30 g/l malt extract, 1 g/l peptone, 15 g/l agar) was used as basis to investigate growth of the three *N. moseri* strains under different stress conditions.

The strains were incubated at different temperatures (37 °C, 28 °C, and 21 °C) in order to narrow down the possible optimal growth temperature of the strains.

For the purpose of testing the tolerance to increasing salinity, NaCl was added to the MEX plates to a final concentration of 0, 0.5, 1, 1.5, 2, and 2.5 M, respectively.

Additionally, the tolerance of the strains to varying pH in the medium was tested. Therefore, the MEX plates were adjusted to pH 2, 3, 5, 7, 8, and 9 with HCl and NaOH, respectively, under sterile conditions. 10g/l Phytagel + 5 mM MgCl_2_ was used instead of agar in those plates. The plates used for salinity- and pH-tolerance-testing were incubated at 28 °C. All plates were inoculated by applying 5 µl of a spore solution (OD_600_ of 3) to the center of the plates and were incubated at the according temperatures for 11 days.

The resulting colony radii were measured using FIJI (ImageJ2, Version 2.9.0).

### BIOLOG assay

Growth of *N. moseri* strains on 95 different carbon sources was performed using the BIOLOG FF (Filamentous Fungi) MicroPlate™ (Art.Nr. 1006) panels (Biolog, Hayward, CA, United States). Spore solution was applied to FF Inoculating Fluid (Art.Nr. 72106) to a turbidity-increase of 20% (Turbidity Deice Name) and incubated for 16 days at 28 °C. Fungal growth was determined by measuring optical density at 750 nm (OD_750_) using a plate reader (TECAN Spark® Multimode Microplate Reader) after each 24 hours, starting after inoculation (day 0). The assay was performed in technical triplicates. We decided to compare OD_750_max independent from growth rate as a simple method to estimate the potential biomass formation on carbon sources. Data was evaluated using GraphPad Prism 9.1.2 (GraphPad Software, LLC.). Statistics were performed using GraphPad Prism 9.1.2 (GraphPad Software, LLC.).

## Acknowledgements

We kindly thank Gerd Mauschitz and Wolfgang Ipsmiller (both TU Wien, Vienna, Austria) for providing access to the scanning electron microscope and supporting us with the measurements. The authors are thankful for the arboreal field work performed by Alexey Kopchinskiy (Austria) in a frame of WWTF-LS13-048 project with kind assistance of the staff from the Kuala Belalong Field Studies Centre (Brunei Darussalam). We thank Kamariah A. Salim and Linda B.L. Lim (Universiti Brunei Darussalam) for their contribution in taxonomic identification of the host plants.

## Author contributions

NJH performed the microbiological assessment, the antiSMASH-analysis, and co-drafted the manuscript

GAV performed the genome sequencing, assembly, annotation, and co-drafted the manuscript

MS performed the phylogenetic analysis

RLM was involved in the study design, supervision, and provided resources ARMA was involved in the study design, supervision, and provided resources MJR isolated the two new strains

CMC provided resources and taxonomic identification of the host plants FC was involved in taxonomic evaluation

ISD was involved in the study design, taxonomic evaluation and and co-drafted the manuscript

CZ was involved in the study design, supervision, provided resources, performed the cblaster analyses, and co-drafted the manuscript

