## Supplementary material for "Genome sequencing and physiological characterization of three *Neoarthrinium moseri* strains": Figures S1 - S10, Tables S1 and S2

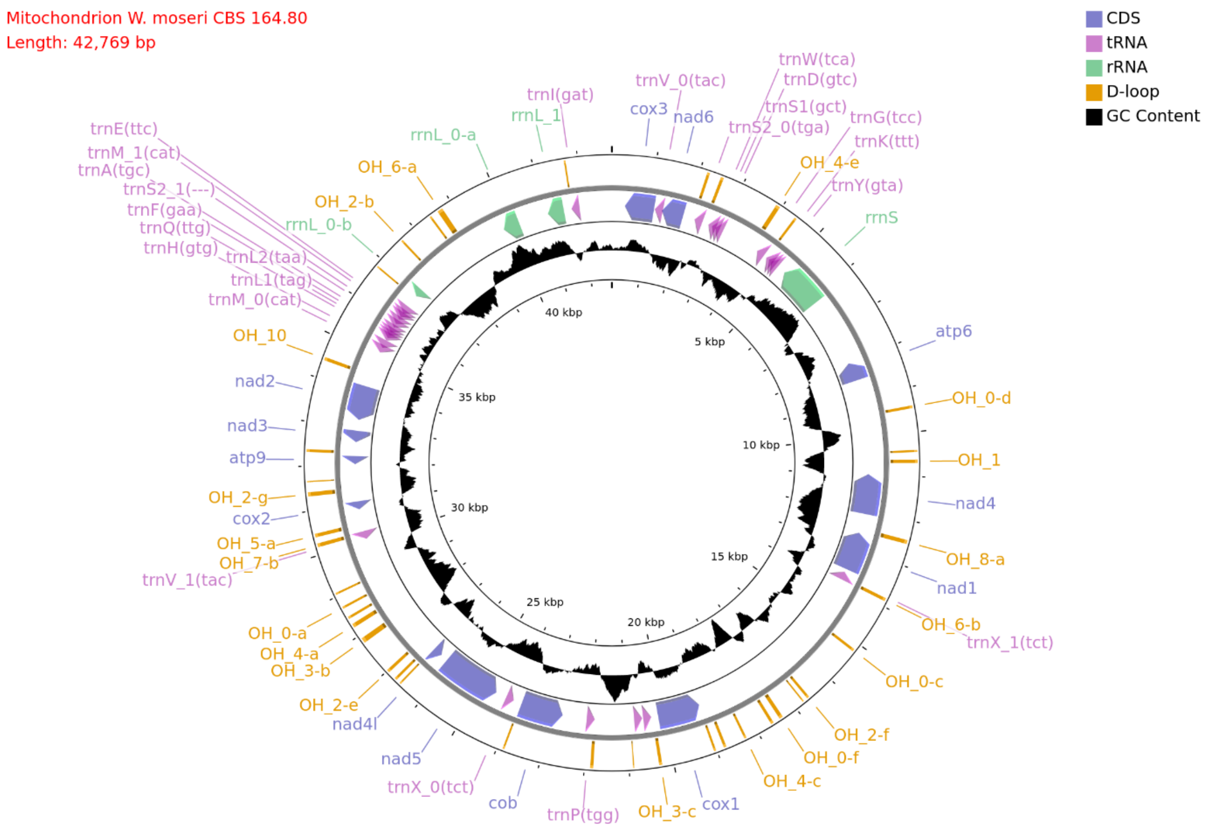


**Figure S1.** Mitochondrial genome of *N. moseri* CBS 164.80.

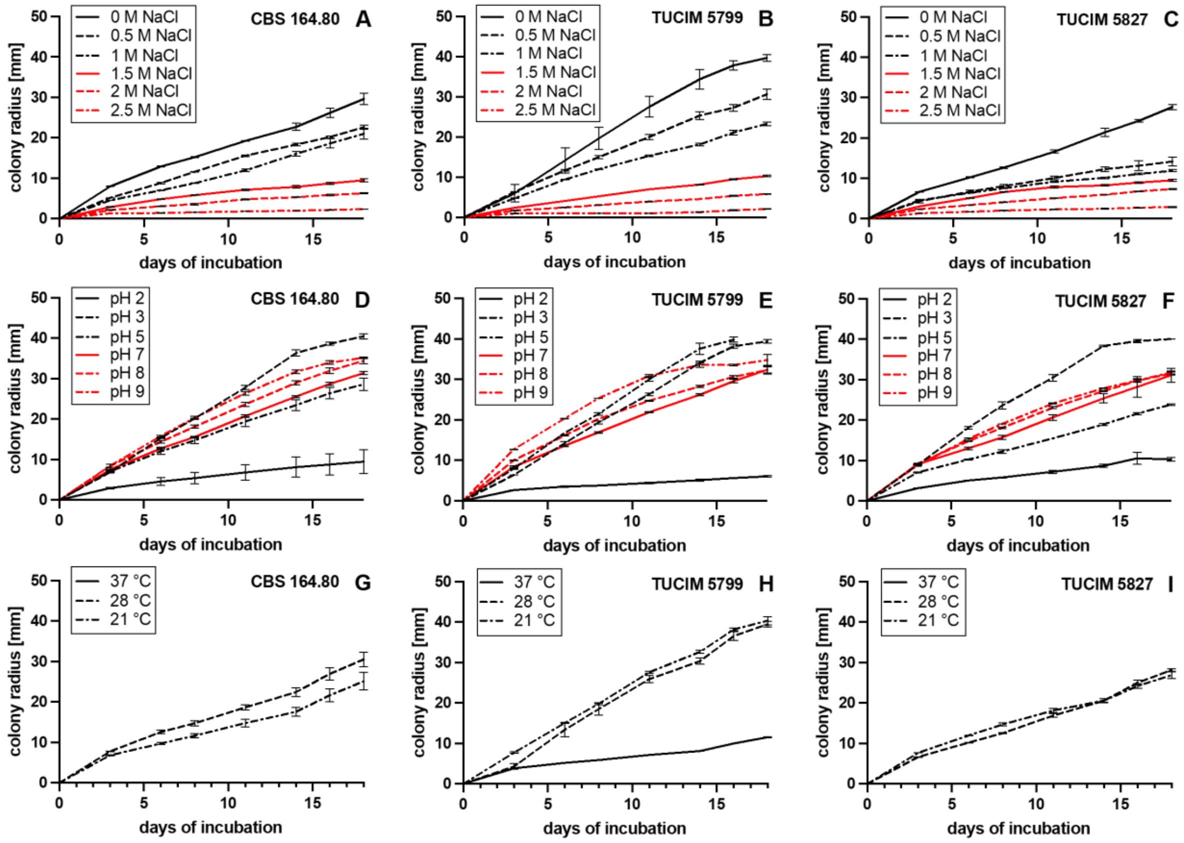


**Figure S2**. Growth of *N. moseri* CBS 164.80 (A/D/G), TUCIM 5799 (B/E/H), and TUCIM 5827 (C/F/I) on medium containing different NaCl-concentrations (A-C), adjusted to different pH (D-F), and incubated at different temperatures (G-I), respectively. Data show mean of three experiments ± SD

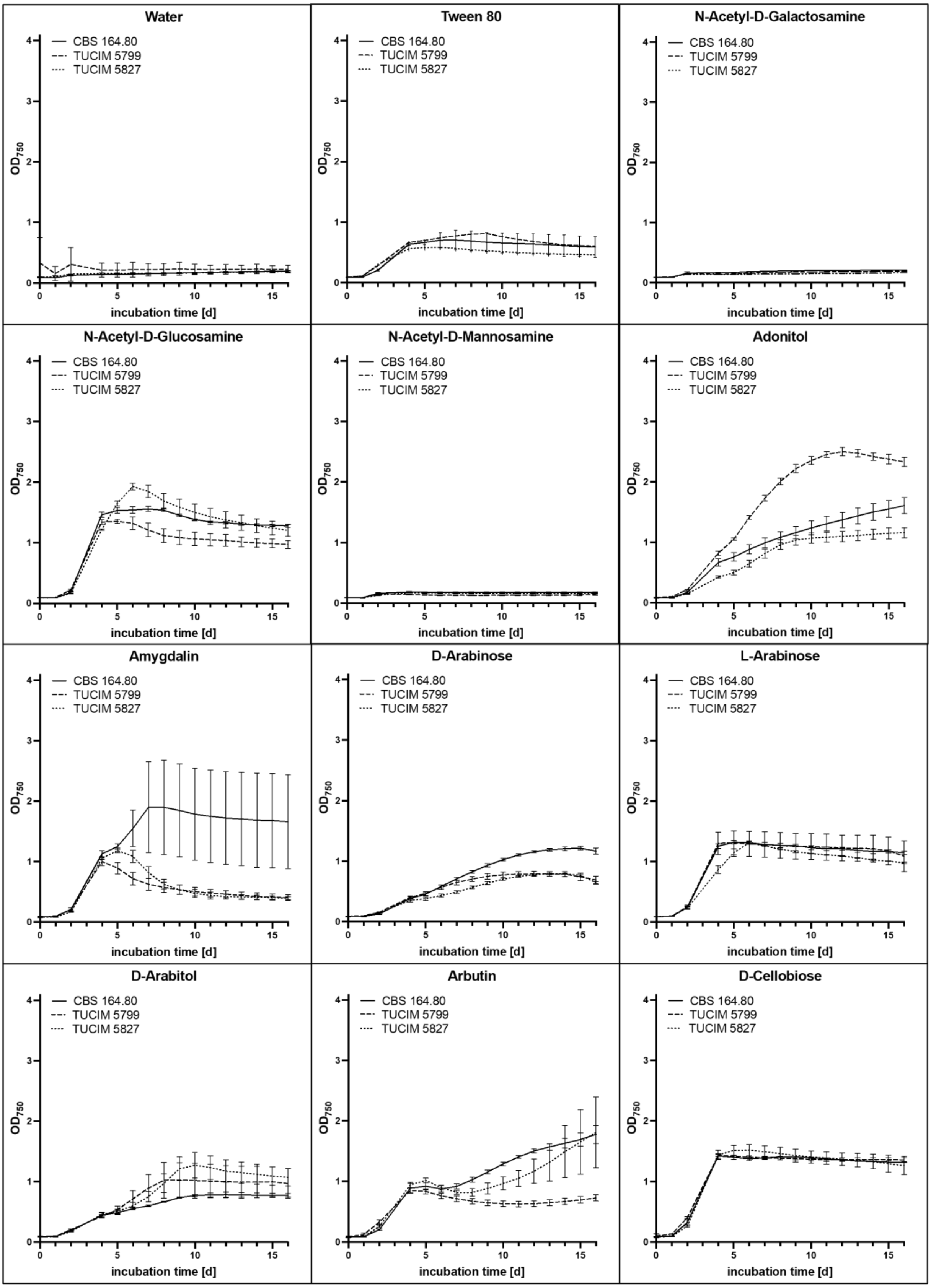

Water
Tween 80
N-Acetyl-D-Galactosamine
N-Acetyl-D-Glucosamine
N-Acetyl-D-Mannosamine
Adonitol
Amygdalin
D-Arabinose
L-Arabinose
D-Arabitol
Arbutin
D-Cellobiose
CBS 164.80
TUCIM 5799
TUCIM 5827
$OD_{750}$
incubation time [d]

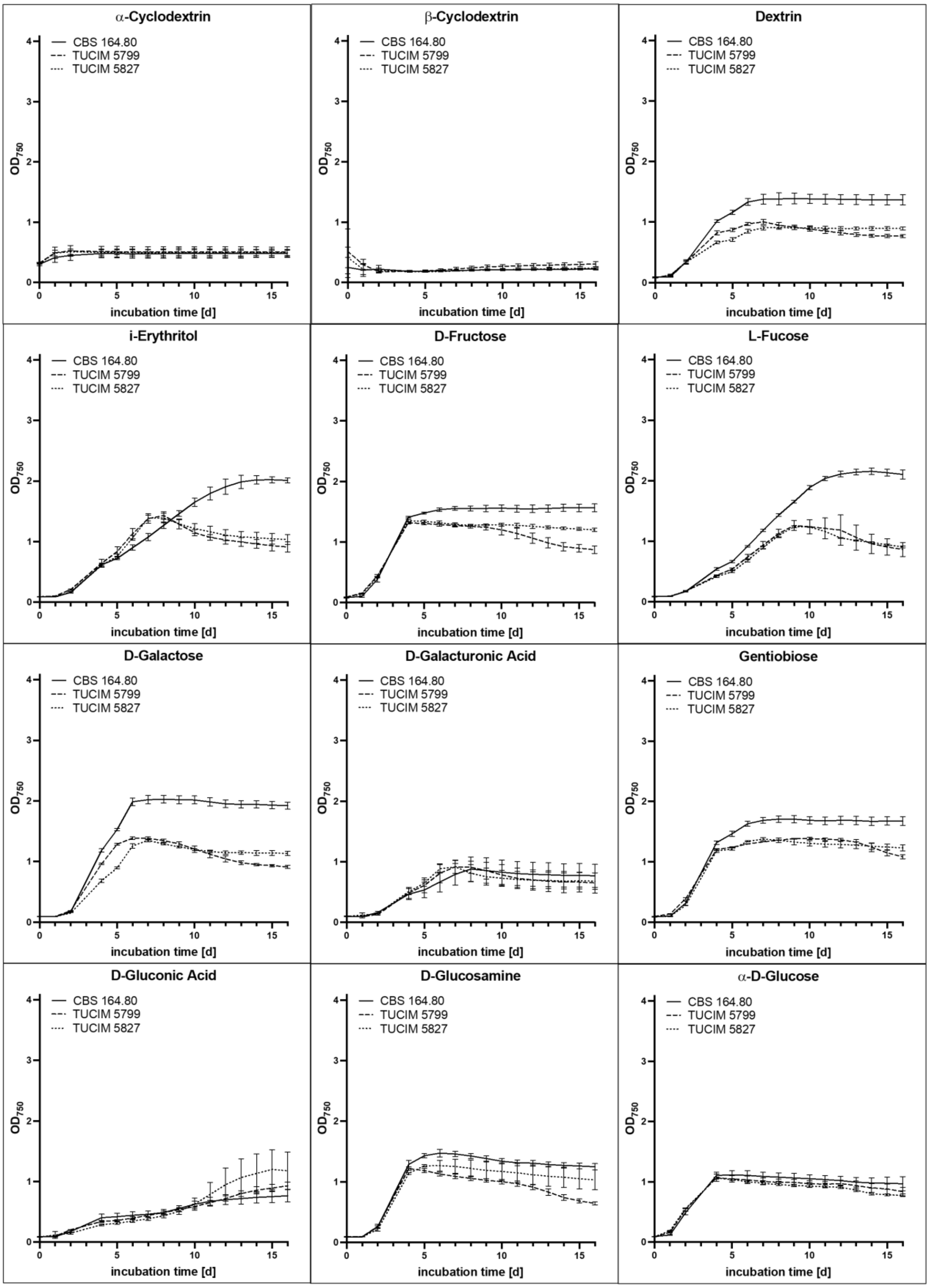


**Figure S4**. Growth of *N. moseri* CBS 164.80 (A/D/G), TUCIM 5799 (B/E/H), and TUCIM 5827 (C/F/I) on different carbon sources in the BIOLOLG assay.

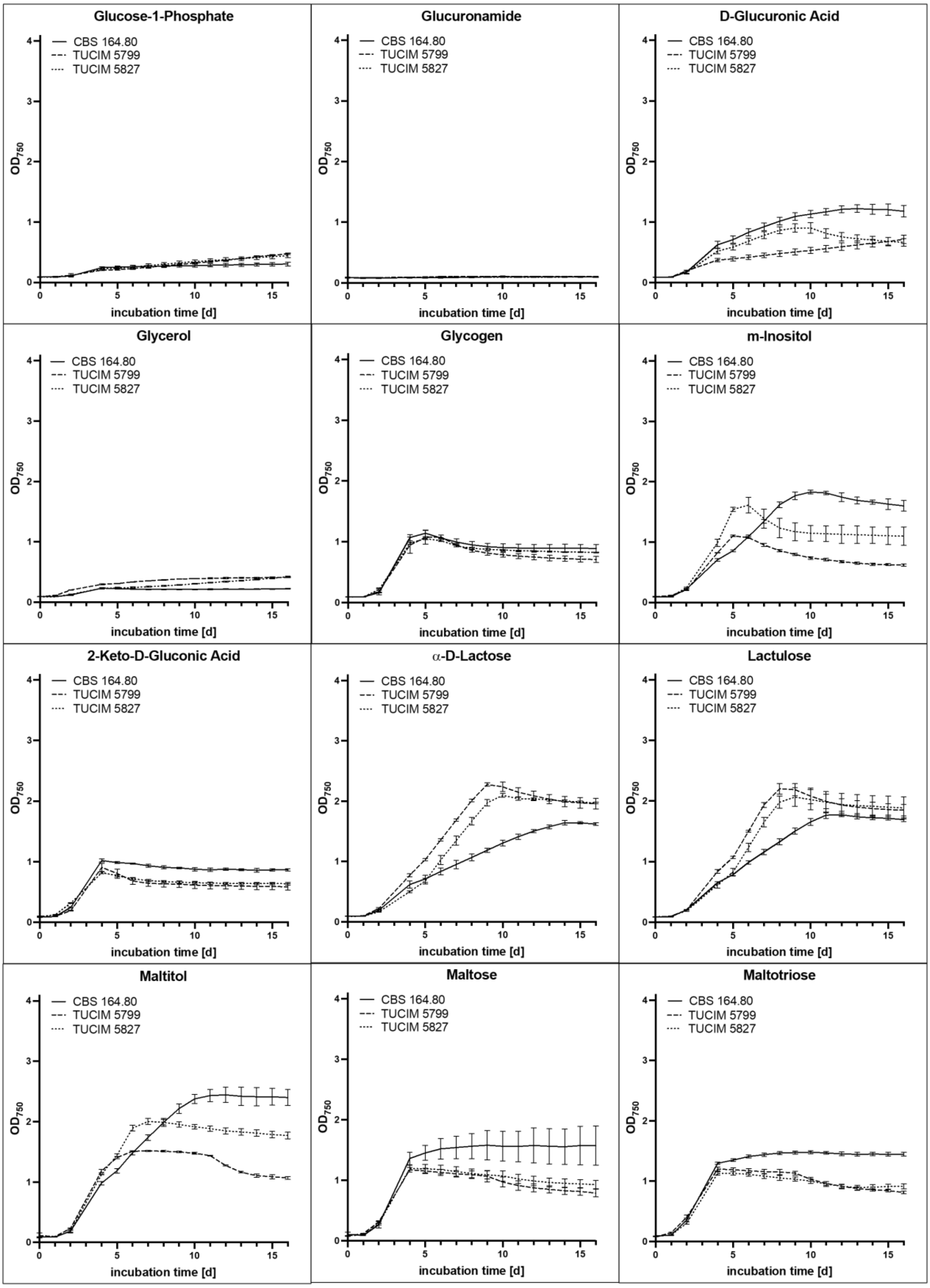


**Figure S5**. Growth of *N. moseri* CBS 164.80 (A/D/G), TUCIM 5799 (B/E/H), and TUCIM 5827 (C/F/I) on different carbon sources in the BIOLOLG assay.

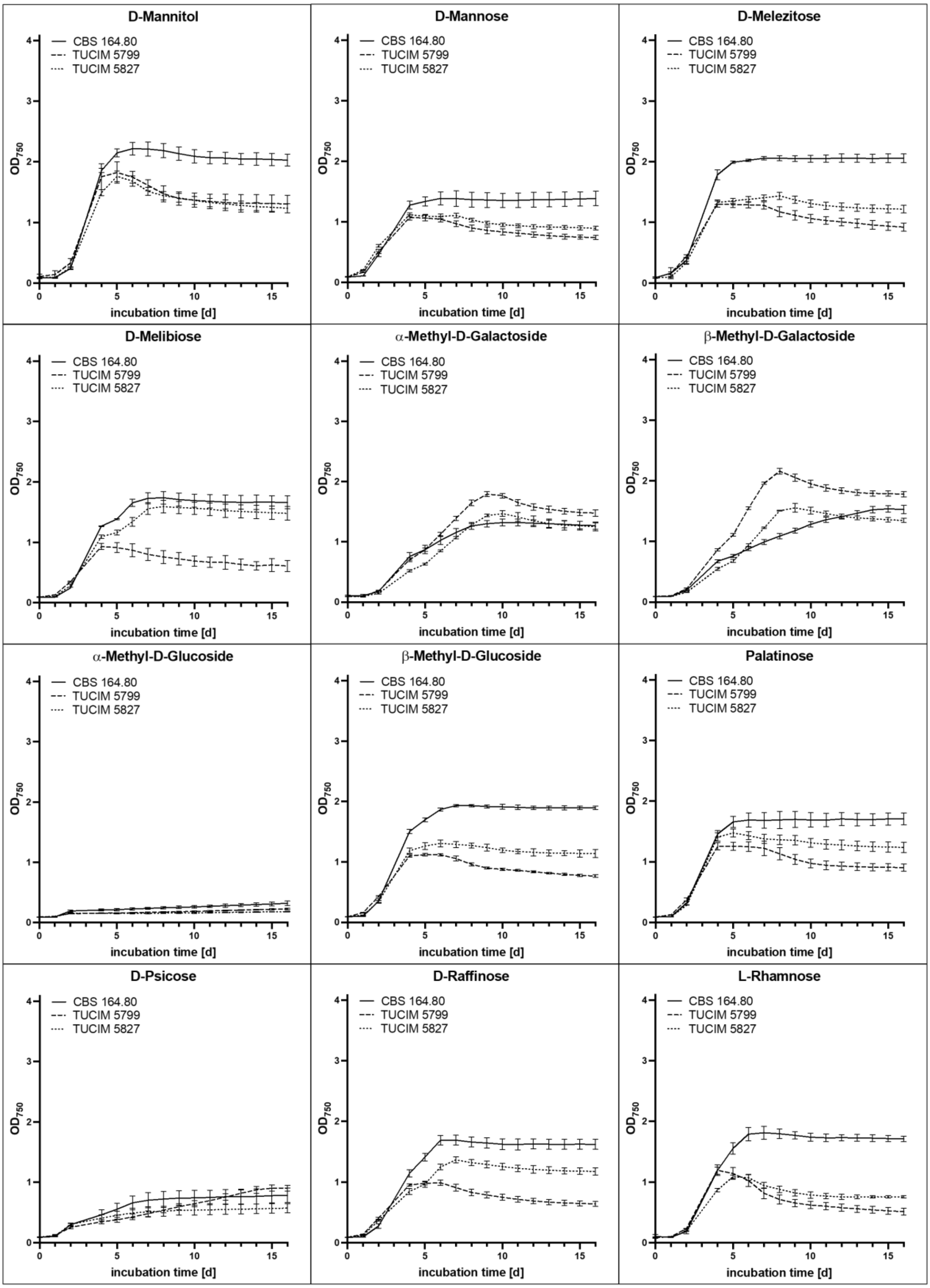


**Figure S6**. Growth of *N. moseri* CBS 164.80 (A/D/G), TUCIM 5799 (B/E/H), and TUCIM 5827 (C/F/I) on different carbon sources in the BIOLOLG assay.

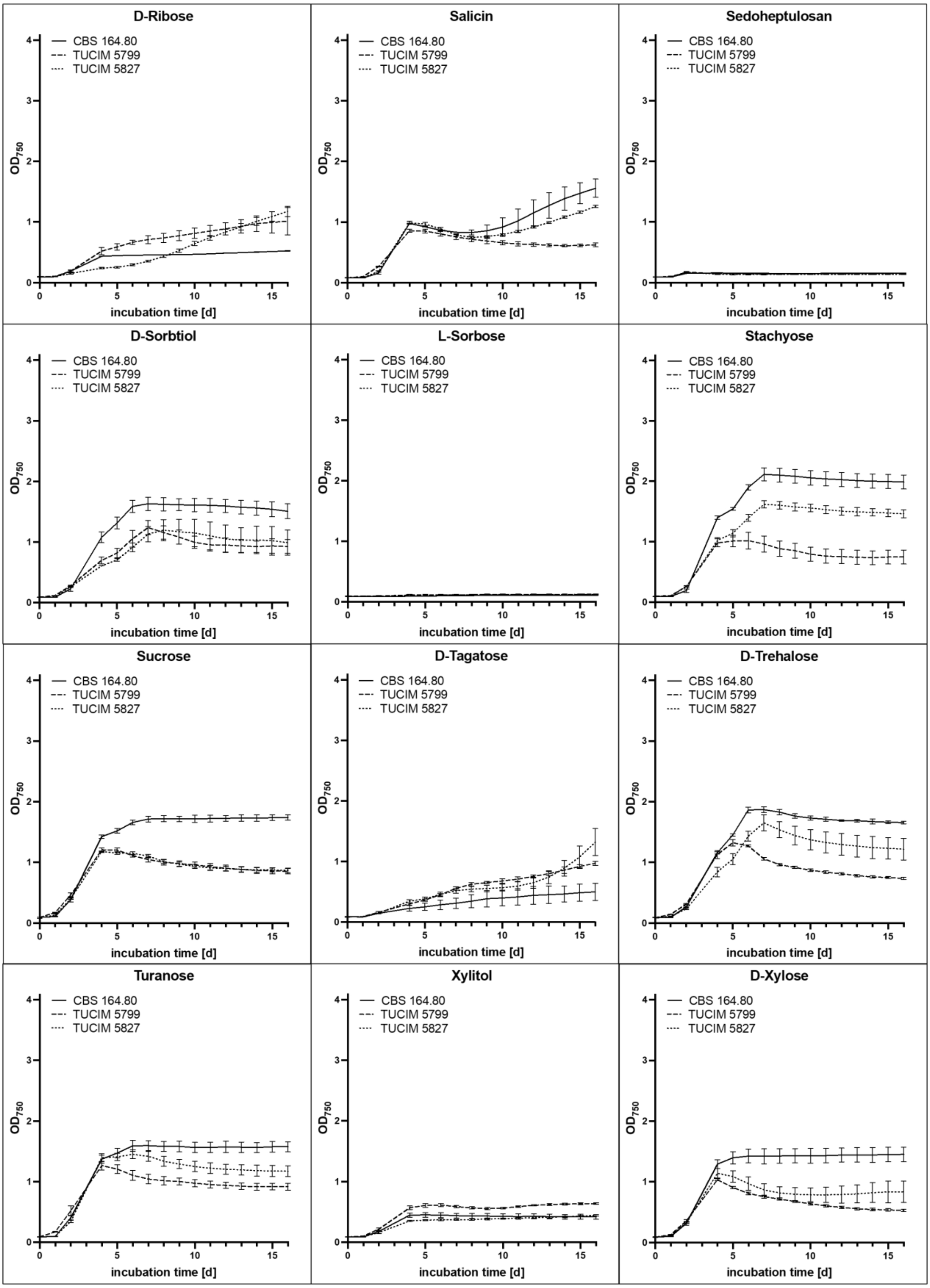


**Figure S7**. Growth of *N. moseri* CBS 164.80 (A/D/G), TUCIM 5799 (B/E/H), and TUCIM 5827 (C/F/I) on different carbon sources in the BIOLOLG assay.

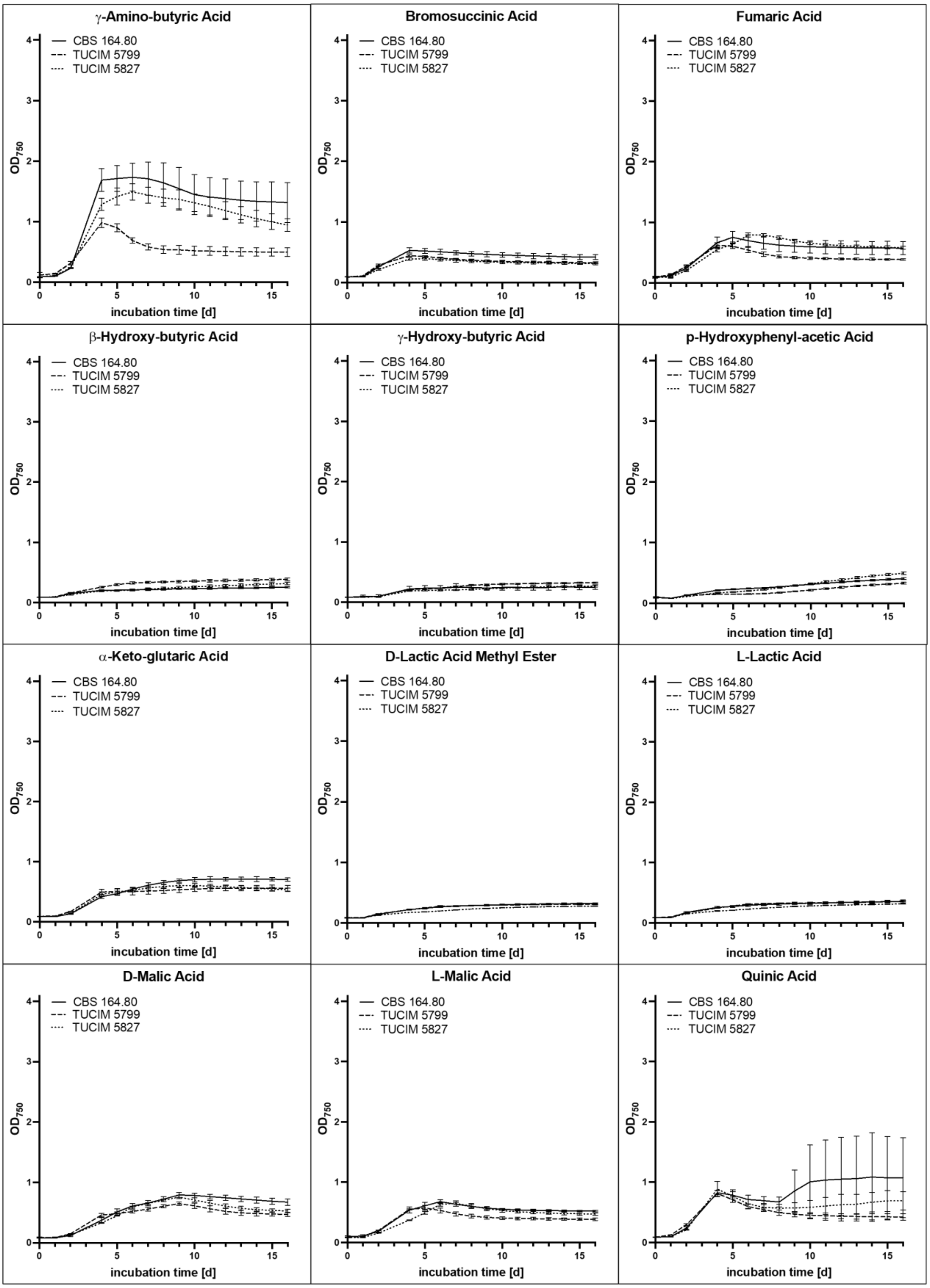


**Figure S8.** Growth of N. moseri CBS 164.80 (A/D/G), TUCIM 5799 (B/E/H), and TUCIM 5827 (C/F/I) on different carbon sources in the BIOLOLG assay.

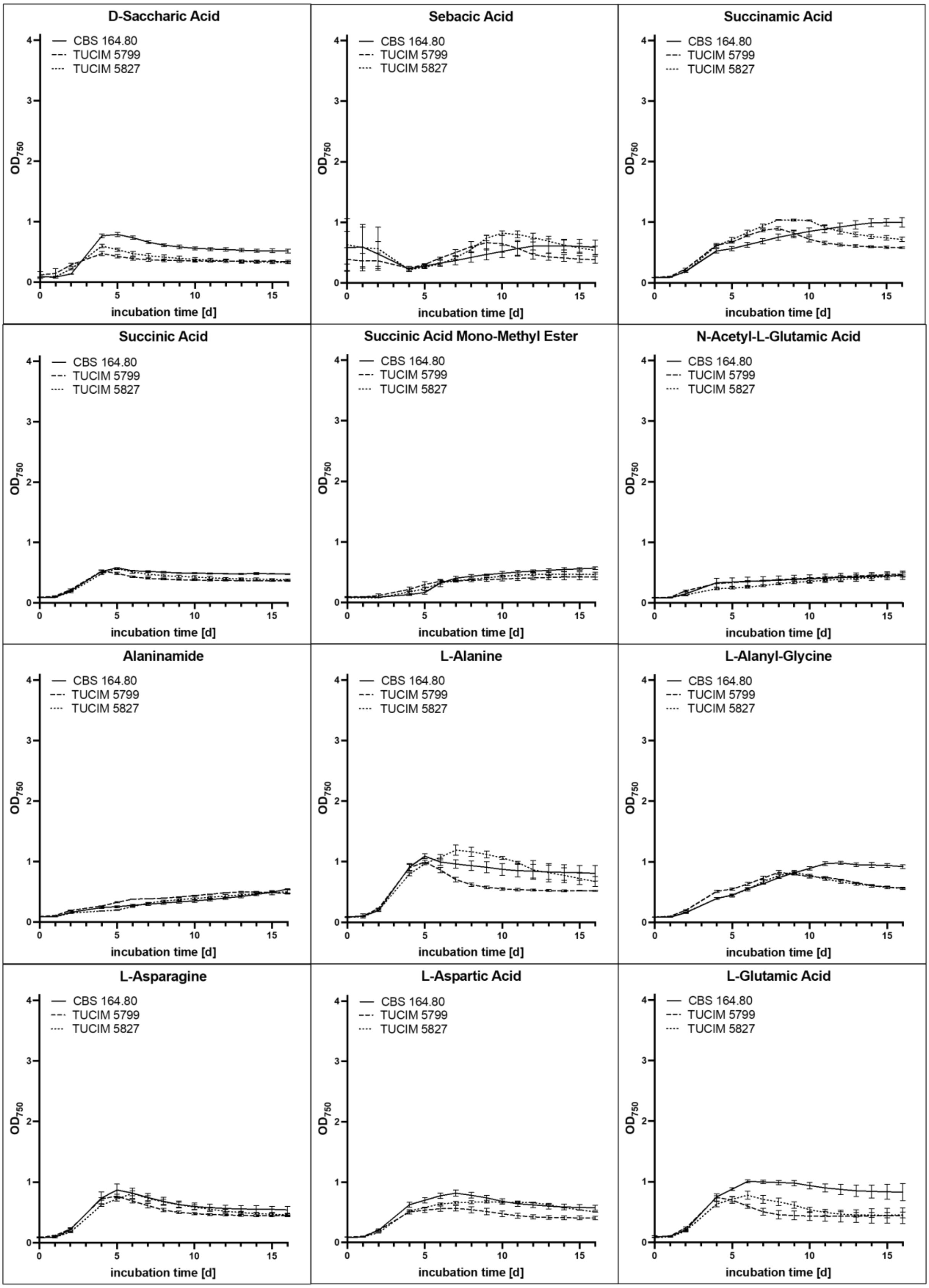


**Figure S9**. Growth of *N. moseri* CBS 164.80 (A/D/G), TUCIM 5799 (B/E/H), and TUCIM 5827 (C/F/I) on different carbon sources in the BIOLOLG assay.

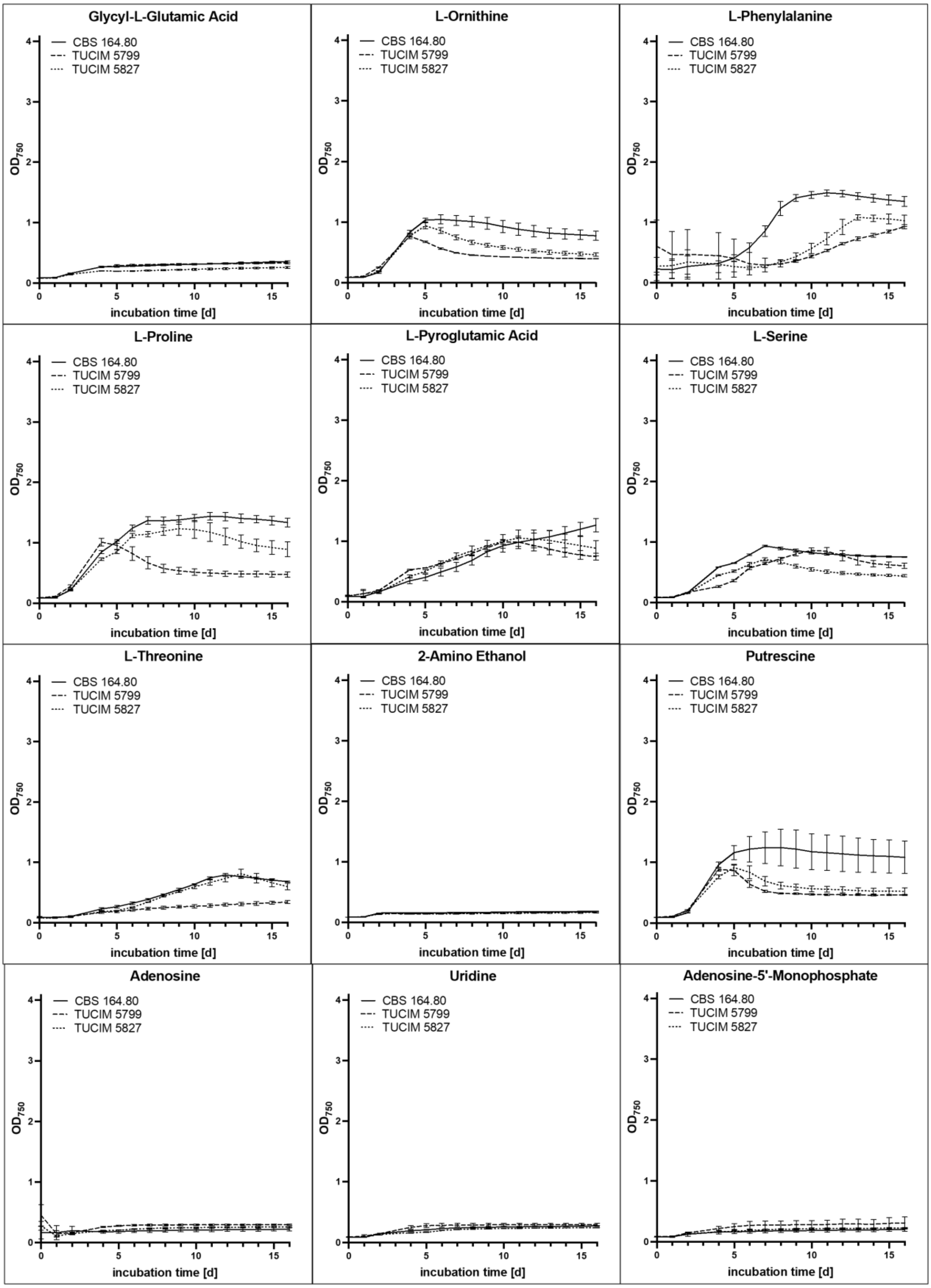


**Figure S10**. Growth of *N. moseri* CBS 164.80 (A/D/G), TUCIM 5799 (B/E/H), and TUCIM 5827 (C/F/I) on different carbon sources in the BIOLOLG assay.

**Table S1. Genome assembly characteristics.**

| Genome | CBS 164.80 | TUCIM 5827 | TUCIM 5799 |
|---|---|---|---|
| Assembly size (bp) | 43,702,215 | 46,154,457 | 44,394,130 |
| G+C content (%) | 52.77 | 52.65 | 52.66 |
| Scaffolds (>= 0 bp) | 230 | 2730 | 693 |
| Scaffolds (>= 1000 bp) | 193 | 609 | 221 |
| Largest scaffold (bp) | 2,337,669 | 1,719,970 | 2,329,648 |
| N50 (bp) | 506,940 | 462,712 | 764,765 |
| L50 (scaffolds) | 26 | 30 | 17 |
| N's per 100 kbp | 2.33 | 2.17 | 1.81 |
| Complete BUSCO (%) | 100.00 | 100.00 | 100.00 |
| Partial BUSCO (%) | 0.00 | 0.00 | 0.00 |

**Table S2. Average nucleotide identity (ANI) between the *N. moseri* strains.**

| Genomes compared | ANI |
|---|---|
| CBS 164.80 : TUCIM 5799 | 99.0276 |
| CBS 164.80 : TUCIM 5827 | 99.0091 |
| TUCIM 5799 : TUCIM 5827 | 99.1092 |
